# Cellular integration of surface and nutrient signals during the early stage of filamentous growth in *Saccharomyces cerevisiae*

**DOI:** 10.64898/2026.09.02.748872

**Authors:** Shashank Pandey, Marine Louvet, Hyojun Kim, Delphine Studer, Valentin Borgeat, Vincent Vincenzetti, Yves Dusserre, Morgan Delarue, Serge Pelet

**Author notes:** Department of Plant Molecular Biology, University of Lausanne, Lausanne, Switzerland (Delphine Studer). Laboratory of Microbiology and Microtechnology, Ecole Polytechnique Fédérale de Lausanne, Lausanne, Switzerland (Valentin Borgeat). Equal contributions (Marine Louvet, Hyojun Kim, Delphine Studer).

## Abstract

Budding yeast cells can transition from vegetative to filamentous growth under specific environmental stimuli. *Saccharomyces cerevisiae* has served as a model to understand this process, which is linked to the virulence of many fungal pathogens. One manifestation of filamentation in *S. cerevisiae* is the formation of pseudohyphal cells, which are elongated and divide in a polarized manner. Here, we established a live-cell imaging assay to monitor this morphogenetic transition and developed a set of fluorescent expression reporters to probe the activity of various signaling pathways during the first 10 hours of this transition. This approach allowed us to identify the stimulus that activates the filamentous growth Mitogen-Activated Protein Kinase (fgMAPK) pathway, which has long evaded identification. Our results indicate that the fgMAPK cascade senses mechanical cues and becomes activated when cells grow on a solid surface. In addition, we have observed that the stimulation of the transcription factor Gln3 regulated by the TOR pathway in low ammonium condition correlates with the elongation of the cells.

## Introduction

Morphogenetic transitions play a central role in the lives of numerous yeast species. These differentiation processes allow cells to alter their growth pattern, cell shape or colony morphology to adapt to their surrounding environment ^1–4^. Notably numerous fungal pathogens are capable of performing these transitions, which help them to adapt to their host by invading into tissues ^5,6^ or contribute to their transmission by attaching to abiotic surfaces ^7,8^.

The formation of biofilms, the invasion into a substrate and the formation of hyphae represent various manifestations of filamentous growth (FG) ^9^. Depending on the fungal species, a different set of environmental cues can induce these transitions. In many instances, the signaling activities of multiple pathways are integrated by the cells to promote the morphogenetic transition ^2,10^. Interestingly, a common set of regulators is generally activated in the various yeasts that undergo filamentation, such as nutrient sensing pathways and Mitogen-Activated Protein Kinase (MAPK) cascades ^4,11^. These pathways will activate a complex gene expression program implicating multiple transcription factors (TF) that contribute to the differentiation of the cells.

Filamentous growth has also been observed in the budding yeast *Saccharomyces cerevisiae* ^4,9^. This model organism can form biofilms, invade into the agar and produce pseudohyphae ^12–14^. Pseudohyphae are characterized by elongated cells that divide in a directed fashion ^15^. This morphogenetic switch is triggered by low nutrients, although additional cues such as low pheromone concentration or small alcohols have been described to stimulate this cell fate decision ^13,16–18^.

Poor or low levels of nitrogen (N) and/or carbon (C) sources will trigger FG. Thus, the TOR (Target of Rapamycin) pathway responsible for nitrogen sensing and the PKA (Protein Kinase A) pathway along with the SNF (Sucrose Non-Fermenting) pathway responding to carbon source availability have all been shown to contribute to this cell fate differentiation ^16,19–21^ (Fig S1). In haploid or diploid cells, when nutrients are plentiful, TOR and PKA are fully active ^22^. When either N- and C-sources are absent, TOR and PKA pathways are inhibited, and cells enter in stationary phase. Between these two extreme cases, low nutrient levels can trigger the FG response. In the TOR pathway, it has been demonstrated that the kinase Tap42 and the phosphatase Sit4 are implicated in FG induction, while the kinase Sch9 does not seem to contribute to this phenotype ^20^. In rich conditions, Gln3 is maintained in the cytosol by TOR1 phosphorylation, but upon N-source limitation, it is dephosphorylated by Sit4 and enriched in the nucleus to drive the transcription of genes required for nitrogen uptake ^23,24^. The TF Gcn4, which is mainly activated by amino acid starvation via Gcn2, can also respond to other stresses, notably TOR inactivation. In the PKA pathway, in the absence of C-source, Bcy1 inhibits the three kinases Tpk1, Tpk2 and Tpk3, allowing the activation of the general stress response TF Msn2 ^29,30^. However, under FG inducing conditions, the kinase Tpk2 is believed to play a specific role in promoting FG via the activation of the TF Flo8 ^31^. In parallel, Tpk1 and Tpk3, when fully active, will negatively regulate the transition to FG ^32^. The SNF pathway is activated in the presence of low glucose or poor C-source. The Snf1 kinase releases the catabolic repression operated by Mig1 and promotes the transcription of gluconeogenesis genes via Cat8 and Sip4 ^33^.

In parallel to these nutrient sensing pathways, the filamentous growth MAPK (fgMAPK) is also required for eliciting the FG phenotype and has been described to be activated in low nutrient conditions ^14,34,35^. Genetic analyses of the fgMAPK cascade have shown that it shares common elements with the mating pathways, such as the MAP kinase kinase kinase (MAP3K) Ste11, the MAP kinase kinase (MAP2K) Ste7 and the MAPK Kss1. Active Kss1 phosphorylates Dig1 and Dig2, relieving the repression on the heterodimer formed by Ste12 and Tec1 ^36,37,34^. The membrane sensors Msb2, Sho1 and Opy2 have been reported to control the activation of the fgMAPK cascade ^38–41^. However, it remains unclear if and how nutrient levels are sensed by these surface proteins and relayed to the MAPK cascade.

It has been established that the transition from vegetative growth (VG) to filamentous growth requires the activation of multiple pathways. Their activity is integrated by the cell to promote the morphogenetic switch ^9^. Interestingly, the inhibition of one pathway can be compensated by the hyper activation of another ^20^. This suggests that there is a complex regulatory network that links these pathways. Although many cross-inhibition or cross-activation mechanisms may exist at the signaling activity level, one well-established layer of signal integration happens at the transcriptional level. Many promoters of the genes expressed upon FG induction harbor binding sites for multiple TFs. One prime example is of this complex regulation is the promoter of the flocculin gene FLO11, which is controlled by PKA, SNF, fgMAPK and amino acid starvation via Gcn4 ^42–45^.

To decipher the contributions of these various signal transduction cascades to the switch from VG to FG, we have developed fluorescent reporters to monitor the dynamics of activation of these pathways under different environmental conditions. We uncover a dynamic interplay between Gcn4 and Gln3 that correlates with the switch to an elongated morphology. In addition, our results indicate that the fgMAPK pathway does not sense nutrient levels but rather responds to mechanical cues from the environment.

## Results

To monitor the switch from vegetative to filamentous growth, we have developed live-cell imaging assays, where cells are growing at the bottom of a well slide embedded in agarose. Prior to the imaging, log-phase growing diploid yeast cells from the Σ1732b background are filtered and washed with YB (Yeast Base, containing essential minerals and vitamins but no N- or C-source) to remove the nutrients present in the growth medium and resuspended in YB (Fig S2). The cells are subsequently loaded into a well and the molten low-melting agarose with defined concentrations of ammonium and glucose is placed on top of the cells. Under these conditions, we can monitor by time-lapse imaging the formation of micro-colonies for up to 12 hours and quantify the induction of fluorescent reporters in the growing cells.

### Flo11 Induction

To test the FG induction in this microscopy assay, we initially set out to develop fluorescent expression reporters based on endogenous promoters that would correlate with FG induction. One obvious candidate was the flocculin Flo11, which contributes to the filamentation phenotype by promoting adhesion to surfaces and cellular aggregation via the A-and B-domains ^46,47^.

Induction of the FLO11 gene has been used in numerous studies as a proxy for FG induction ^32,42,44,48^. The 3.2 kB sequence of the *FLO11* promoter was cloned in front of a quadruple-Venus (qV) reporter construct and integrated into the *URA3* locus using a single integration vector ^49^ (Fig 1A). This long promoter includes the regulation by two non-coding RNA (ncRNA) which play a key role in the bimodal induction of the *FLO11* locus ^50,51^. At the onset of the time-lapse, a fraction of cells are already fluorescent. As cells start to grow in Synthetic Low Ammonium Dextrose (SLAD), the number of fluorescent cells and their intensity increase. A peak in the average fluorescence of the population is observed around 6h after the start of the imaging (Fig 1B and C, Supplementary Movie 1). The division pattern and the increase in the eccentricity of the cells denote a switch to a pseudohyphal growth. However, we fail to observe a correlation between the level of p*FLO11*-qV expression and the eccentricity of the cells at the population level across time or at the single cell level at a specific time point (Fig 1D).

**Figure 1.**
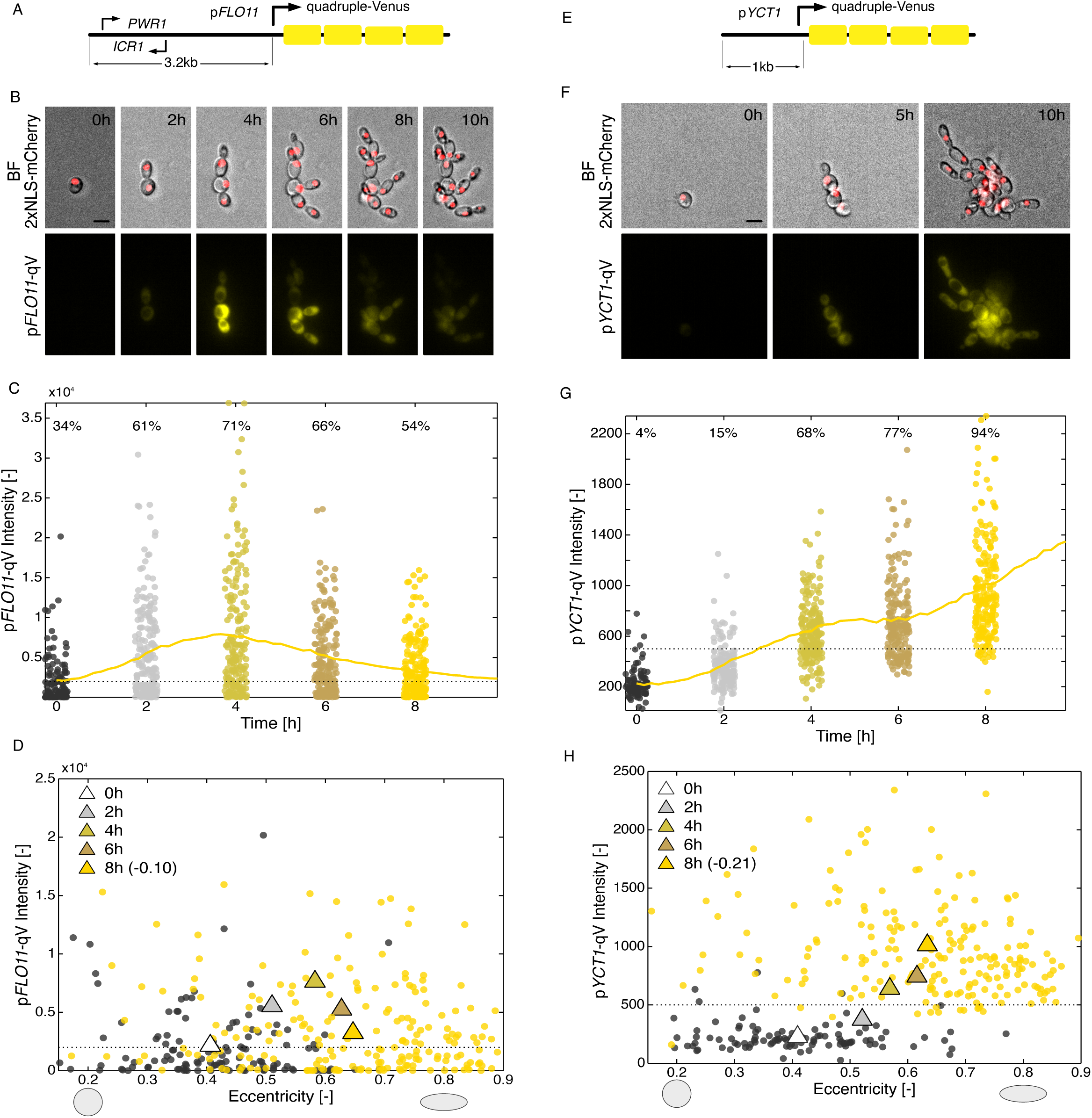
Endogenous promoter for filamentous growth fluorescent reporters. A. The 3.2 kB promoter of *FLO11* is used to control the expression of a quadruple-Venus (qV) construct integrated in the genome. The transcription of the *FLO11* locus is regulated by two long non-coding RNAs (*ICR1* and *PWR1*), which lead to a bistable expression of the locus. B. Time-lapse images of the development of a microcolony of cells bearing the p*FLO11*-qV construct and a nuclear fluorescent tag in SLAD agarose medium. The scale bar represents 5µm. C. Quantification of the p*FLO11*-qV fluorescence signals in cells grown in SLAD. The dots represent the average cellular intensity of individual cells and the solid line represents the mean of the population. The percentage of expressing cells in the population is indicated. The threshold for expression is depicted by the dashed line. For clarity, up to 200 randomly selected cells are displayed. D. Correlation between the eccentricity of the cells and the p*FLO11*-qV intensity. The triangles represent the average eccentricity and intensity of the population at specific time points. The round dots represent individual cell measurements at 0h (black) and 8h (yellow). The gray shapes depict ellipses with an eccentricity of 0.2 (left) and 0.8 (right). For clarity, up to 200 randomly selected cells are displayed. The correlation coefficient between the eccentricity and the fluorescence intensity between all cells measured at 8h is indicated in parentheses. E. Scheme of the p*YCT1*-quadruple-Venus (qV) reporter construct. F. Bright field and fluorescence images of cells bearing the p*YCT1*-qV reporter and growing in SLAD agar environment. The scale bar represents 5µm. G. Quantification of the p*YCT1*-qV fluorescence signals in cells grown in SLAD. The dots represent the average cellular intensity of individual cells and the solid line represents the mean of the population. The percentage of expressing cells in the population is indicated. The threshold for expression is depicted by the dashed line. For clarity, up to 200 randomly selected cells are displayed. H. Correlation between the eccentricity of the cells and the p*YCT1*-qV intensity. The triangles represent the average eccentricity and intensity of the population at specific time points. The dots represent the correlation of these two measures in individual cells at 0h (black) and 8h (yellow). The gray shapes depict ellipses with an eccentricity of 0.2 (left) and 0.8 (right). For clarity, up to 200 individual cells are displayed. The correlation coefficient between the eccentricity and the fluorescence intensity between all cells measured at 8h is indicated in parentheses.

To verify if the two endogenous loci of *FLO11* were regulated similarly to our expression reporter, we inserted a fluorescent protein into the *FLO11* sequence between the A and B domains (Fig S3A). Since both alleles were tagged with different fluorophores (mCherry and mCitrine), we can observe the expression from individual alleles (Fig S3B, Supplementary Movie 2). Unfortunately, due to different properties of the fluorophores, the dynamics of fluorescence apparition are not similar for both alleles (Fig S3C and D). In addition, each allele can be differentially activated in each cell (Fig S3E).

Previous studies have observed a correlation between *FLO11* expression levels and FG. These measurements were obtained using population-level assays and were usually performed at a much later time after FG induction ^42,48^. Under our experimental conditions with single cell resolution, the bistable regulation of the gene leads to a complex expression pattern with cells inducing the construct in VG and a heterogenous apparition of the fluorescence upon FG- inducing conditions that do not coincide with the modification of the cellular morphology.

Therefore, the expression of *FLO11* does not seem to provide a useful proxy for FG induction for single cell studies.

### Endogenous Promoters

The induction of FG coincides with the differential regulation of close to 900 genes ^52^. Based on these transcriptomics data, we sought to identify highly induced genes that could provide a robust readout of FG induction. We cloned the promoters of a few candidate genes in front of the qV reporter and quantified their induction in SLAD conditions. The best response was obtained by the promoter of *YCT1,* which is strongly induced in SLAD medium (Fig 1E and F, Supplementary Movie 3). The fluorescence increases throughout the time-lapse in a homogenous fashion within the population (Fig 1G). At the population level, we observe a combined increase in fluorescence and in the eccentricity of the cells as a function of time. However, this correlation does not hold true at the single cell level, where we observe a negative correlation (-0.21) between eccentricity and cellular fluorescence at the 8h time point (Fig 1H).

Like *FLO11*, *YCT1* is regulated by multiple TFs. According to Yeastract ^53^, the *YCT1* promoter has documented or potential binding sites for Gln3, Gcn4, Msn2/4, Flo8, Cat8, Ste12 and Tec1, among many other TFs. Therefore, the *pYCT1*-qV reporter can potentially integrate signals from TOR, PKA, SNF and fgMAPK pathways. This integration capacity makes this reporter a useful tool to monitor the switch to FG and provides a more straightforward readout than *FLO11* in single cells.

### Synthetic Promoters

The capacity of the *YCT1* promoter to integrate signals from numerous signaling cascades is shared by a large fraction of endogenous promoters, which contain binding sites for multiple TFs. To disentangle the contributions of the different signaling pathways that contribute to the FG response, we decided to engineer synthetic promoters based on the *CYC1* promoting sequence where specific binding sites for individual TF downstream of the TOR, PKA, SNF and fgMAPK pathways were inserted (Fig 2A). For each TF, we engineered promoters with binding motifs based on the known consensus sequences of the TF (pBS) and in parallel a control construct with mutated binding sites (pNB, Fig 2B, Supplementary File 1 (list of TF, binding and non-binding sites)). For Flo8 and Ste12-Tec1, engineering fully synthetic sequences did not result in functional promoters. Therefore, we used longer sequences extracted from endogenous promoters, which have been described to bind specifically these TFs. For Flo8, a 250 bp long sequence extracted from the promoter FLO11 was used ^54^ and for Ste12/Tec1, a 90bp fragment from the Ty1 transposon was selected ^55^. In both instances, a control promoter where a few key bases were mutated was generated to confirm the specificity of the induction.

**Figure 2.**
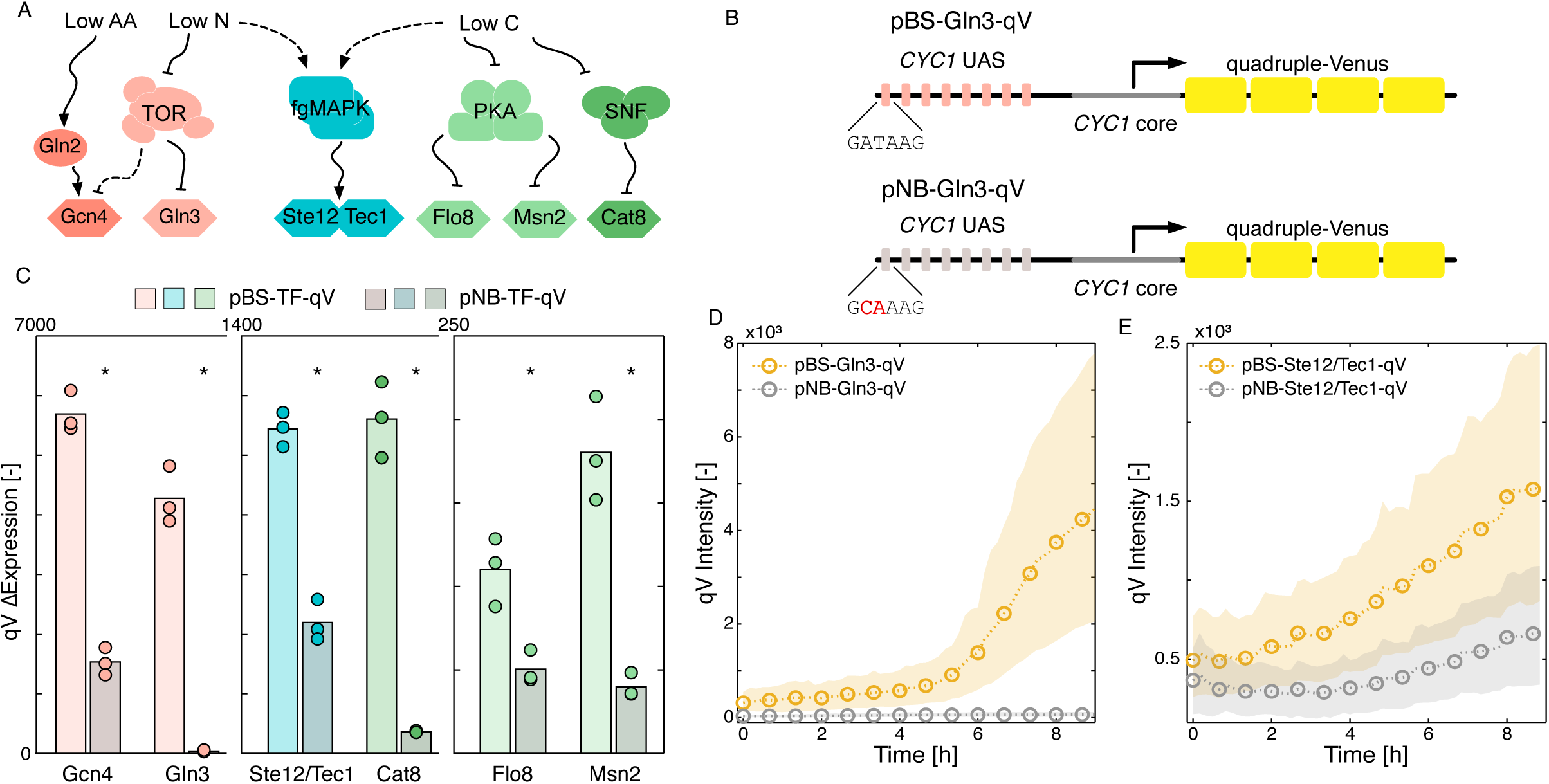
Engineering of synthetic expression reporters. A. Scheme of the signal transduction pathways implicated in FG and the different TFs used to probe the activity in these pathways. B. Description of the generation of the synthetic Gln3 expression reporter, where 7 binding sites (BS) are inserted in the Upstream Activating Sequence (UAS) of the p*CYC1* promoter. A control non-binding (NB) construct is also generated where key residues in the TF-binding site are mutated. C. Differential expression of the synthetic reporters for the six transcription factor dependent reporters, for cells grown in SLAD agar. The bar represents the mean of 3 to 4 biological replicates (indicated by the round markers). The star indicates that the pBS construct is expressed at a significantly different level than the pNB construct (t-test, p-value <0.05). D and E. Dynamics of fluorescence apparition from the binding (yellow) and non-binding (gray) Gln3 (D) and Ste12/Tec1 (E) reporters in cells grown in SLAD agar. The dashed line represents the median of the population and the shaded area, the 10 and 90 percentiles.

Each promoter was cloned in front of the qV fluorescent reporter and integrated into the yeast genome. The expression outputs of the promoters were measured in SLAD agar (Fig 2 C, D and E and Fig S4). In parallel, the constructs were also tested under various stress conditions, such as rapamycin, hyper-osmotic stress, starvation or growth in glycerol (Fig. S5). A significantly stronger fluorescence induction was observed for all the binding compared to the non-binding constructs. In the best cases, the fluorescent induction from the promoter with binding sites was very high, while cells bearing the non-binding construct were non-fluorescent (Fig 2D). In some instances, the non-binding promoters still displayed some level of induction (for Ste12/Tec1 for instance, Fig 2E), suggesting that the few point mutations inserted to decrease the affinity of the TF to the promoter were not sufficient to completely abrogate the binding of the TF to the promoter. None the less, this synthetic promoter strategy allows us to determine which TF is active under the tested environmental conditions and extrapolate the results to determine if or when a signaling cascade is implicated in FG induction.

### Environmental Stimuli

To test the response of the TF to nutrient levels, we defined three nutrient-limiting conditions which could potentially trigger FG and compared them to a rich growth medium (Fig 3A, Supplementary Movies 4, 5, 6, 7). In high glucose (2%) and high ammonium (20 mM) conditions (SAD, Synthetic Ammonium Dextrose), cells maintain their cell size and round morphology throughout the 9 hours of growth (Fig3B and Fig S6A). The microcolonies develop in a cluster of aggregated cells. When the glucose concentration is lowered to 0.05% (SALD, Synthetic Ammonium Low Dextrose), the average cell size drops during the first 2 hours of growth and, in parallel, the mean eccentricity of the population increases steadily from 0.4 to overcome 0.55. When only the ammonium concentration is decreased to 50µM (SLAD), the eccentricity regularly increases up to 6 hours to overcome 0.55, similar to what is observed in SALD. Then, around 6 hours of growth, a second phase of eccentricity increase can be monitored to overcome 0.65 in SLAD conditions. When both ammonium and glucose are limiting (SLALD, Synthetic Low Ammonium Low Dextrose), the rise in eccentricity is faster than if only the ammonium is limiting. However, in parallel, the area of the cells drops and, overall, cells lose close to 40% of their original size. In all cases of nutrient deprivation, the microcolonies shapes are more structured with cells extending from the center of the colony (Fig 3A).

**Figure 3.**
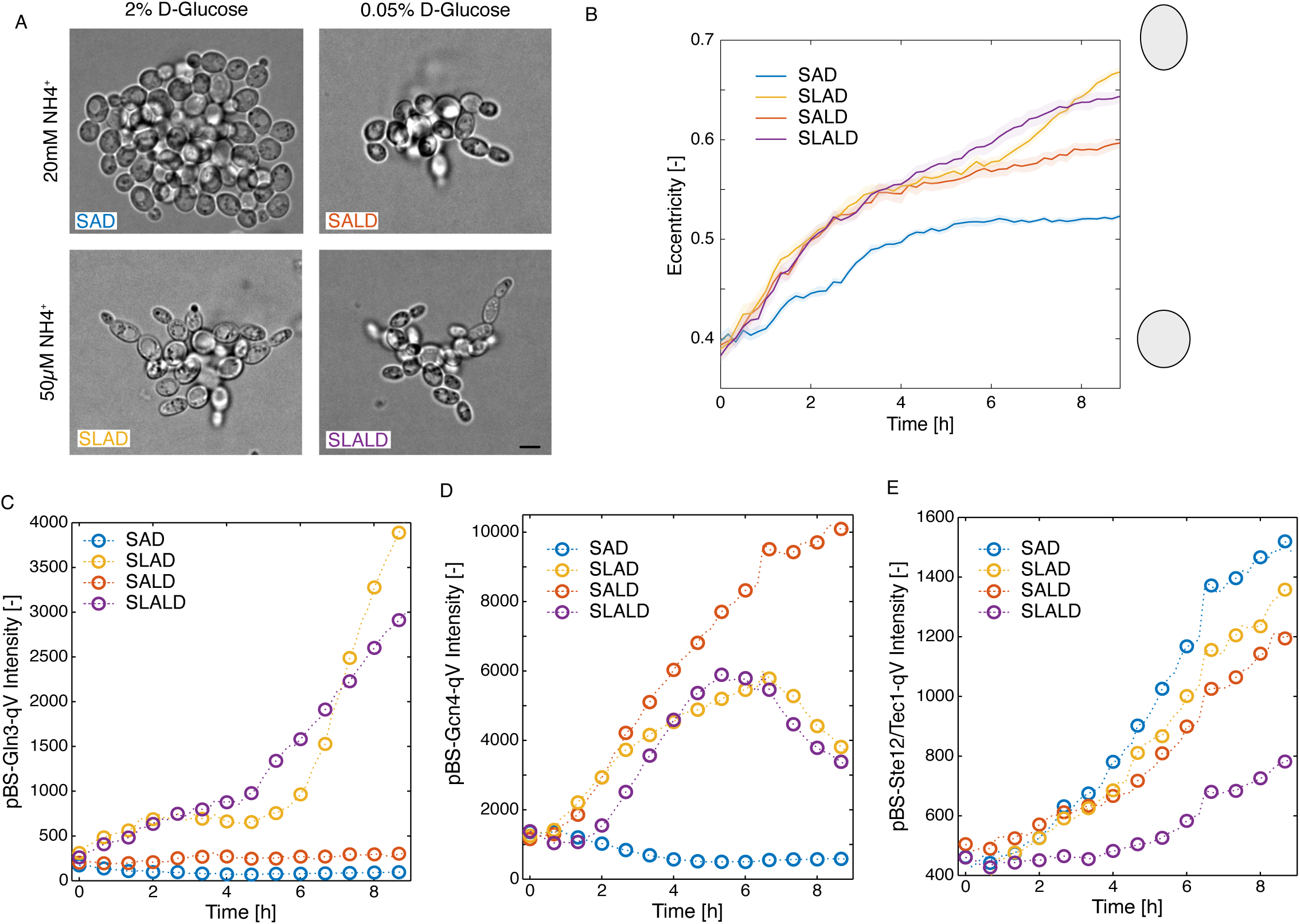
Effect of nutrients on morphology and gene expression. A. Effect of ammonium and glucose concentrations on the cellular morphology and microcolony structure after 9 hours of growth in agar medium. The scale bar represents 5µm. B. Evolution of cellular eccentricity during growth in different media. The solid line represents the mean of 9 population averages obtained from the three biological replicates performed with three different strains. The shaded area represents the standard error of the mean. The gray ellipses drawn on the right represent the shape of an ellipse with an eccentricity of 0.4 (bottom) or 0.7 (top). C, D and E. Dynamics of the fluorescence intensity generated by the qV construct under the control of the synthetic Gln3 (C), Gcn4 (D) and Ste12/Tec1 (E) dependent reporters in cells grown in four different media conditions. The dashed line and markers represent the mean of the population of all the cells measured at each time point.

These four conditions were used to test the induction of our synthetic promoters. For the Msn2- and Cat8 dependent promoters (Fig S6B and C), induction was strongest in the low ammonium medium (SLAD), while it was very limited in low glucose medium (SALD). Weak induction was observed for the pBS-Flo8-qV reporter in all four conditions tested (Fig S6D). The low level of response observed for these three reporters contrasts with the noticeable induction seen in other stimulating conditions (Fig S5 E, F and G). This indicates that these TFs are not highly activated during these first 10 hours of FG induction, suggesting that PKA and SNF are not sufficiently active to produce a sustained transcriptional response in our experimental conditions.

In contrast, a strong fluorescent signal is generated by the Gln3 and the Gcn4-dependent reporters. pBS-Gln3-qV is activated in both media where ammonium is limiting (Fig 3C) in-line with the expected direct connection with the repression of TOR activity in Nitrogen starvation. Gcn4 is activated under all nutrient limiting conditions (Fig 3D) in agreement with its role as a master regulator for starvation signals ^26^. Finally, the reporter for the fgMAPK pathway, based on Ste12 and Tec1 binding sites, is induced in all four conditions, including in rich nutrient environments (Fig 3E). Note that, even for these three synthetic reporters, which are induced in the SLAD conditions, we do not observe a positive correlation between the fluorescence intensity of the reporter and the eccentricity of the cells at the single cell level (Fig S6 E, F, G). Together, these results imply that the downregulation of TOR and the activation of the fgMAPK pathway play a central role in the early phase of FG induction that we probe with our time-lapse assays.

### Interplay Between Gln3 and Gcn4

Interestingly, while the TF Gln3 and Gcn4 are both targets of the TOR pathway, they display drastically different induction dynamics in SLAD, suggesting distinct regulatory mechanisms. Expression from the GCN4-controlled promoter rises rapidly after the transfer to the low ammonium medium, peaks at 6h and then decreases. In contrast, Gln3-driven expression remains low and until it rises suddenly after the 6th hour. Intriguingly, the rise in eccentricity of the cells seems to happen at a similar time point (Fig 3B).

To investigate if there is a temporal link between the Gcn4 and Gln3 activities, a strain combining the pBS-Gln3-qV reporter and a pBS-Gcn4 promoter controlling a triplet-Scarlet (tSc, red) was constructed (Fig 4A, Supplementary Movie 8). The same dynamic interplay could be observed in this dual reporter strain with a transient Gcn4-driven fluorescence induction and a delayed apparition of the Gln3-dependent reporter.

**Figure 4.**
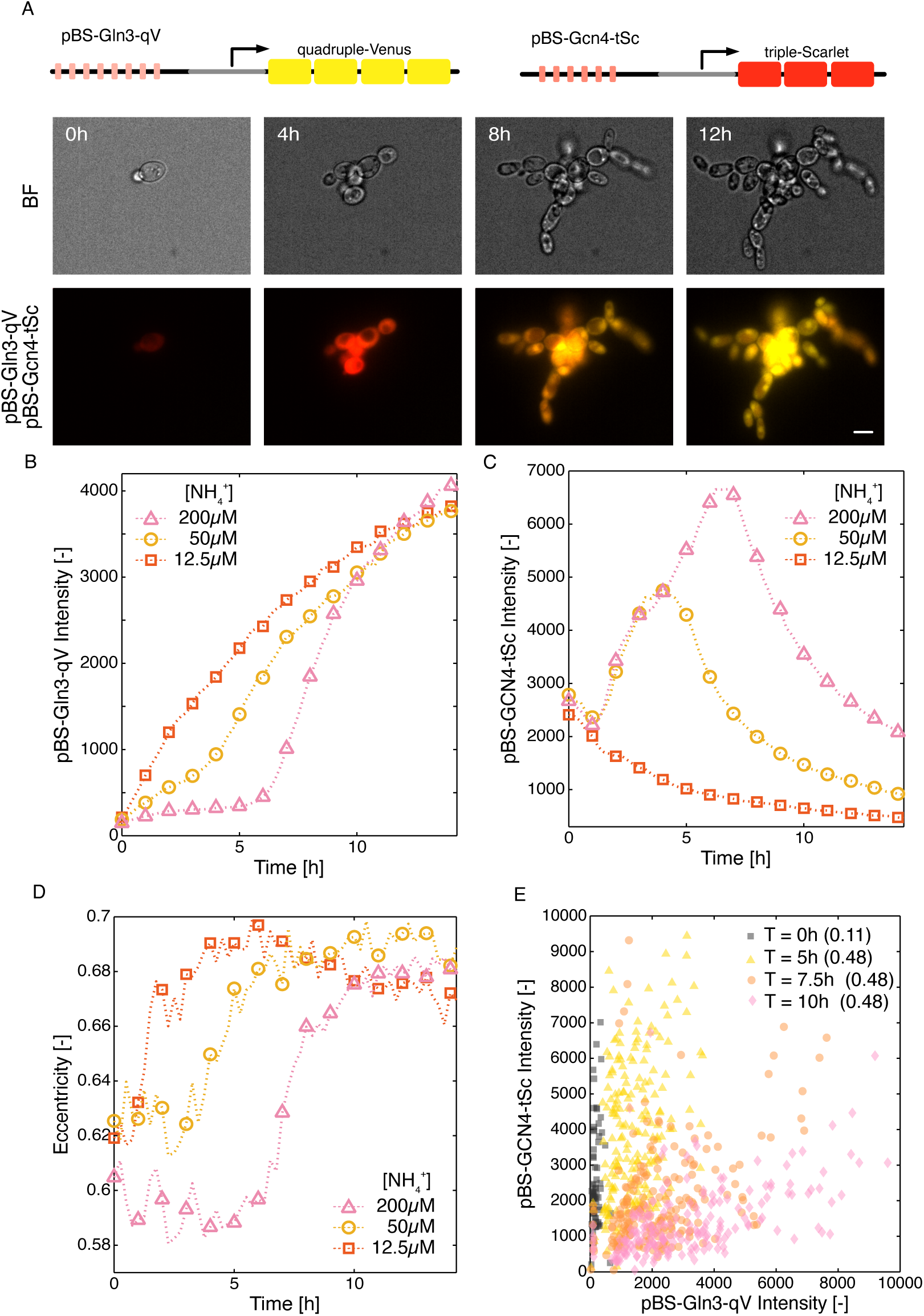
Interplay between Gln3 and Gcn4 activity. A. Schematic of the fluorescent reporters and images of cells containing the pBS-Gln3-qV (yellow) and the pBS-Gcn4-tSc (red) reporters grown in SLAD agarose medium. The scale bar represents 5µm. B and C. Dynamics of the fluorescence intensity generated by the pBS-Gln3-qV (B) and the pBS-Gcn4-tSc reporters in cells grow in SLAD agar medium (round yellow, 50µM NH_4_^+^), four times lower ammonium (square orange, 12.5µM NH_4_^+^) or four times higher ammonium concentration (triangle pink, 200µM NH4^+^). The mean fluorescence measured from one representative replicate is plotted. D. Evolution of the mean cellular eccentricity of the cells grown in SLAD agar medium (round yellow, 50 µM NH_4_^+^), four times lower ammonium (square orange, 12.5 µM NH_4_^+^) or four times higher ammonium concentration (triangle pink, 200µM NH_4_^+^). E. Correlation between the average cellular intensity in the yellow (pBS-Gln3-qV) and red (pBS- Gcn4-tSc) channels after 0, 5, 7.5 and 10 hours growth in SLAD agar medium with 50µM NH_4_^+^. The correlation between the yellow and red fluorescent signal calculated between the measured cells is indicated in parentheses. For clarity, up to 200 randomly selected cells are displayed.

More interestingly, by tuning the concentration of ammonium in the medium at the onset of the experiment, we could determine that the transition between Gcn4 and the Gln3 expression regimes was controlled by the ammonium levels (Fig 4B and C). A higher level of NH_4_^+^

(200µM) delays the Gln3-dependent expression and the peak in pBS-Gcn4 reporter induction. While in low ammonium (12.5µM), the Gln3-dependent expression takes place from the beginning of the experiment, and the Gcn4-driven reporter fails to be induced. Strikingly, the rise in eccentricity in all three conditions is temporally correlated with the induction of the pBS- Gln3-qV construct (Fig 4D). Note that, at the single cell level, we observe a correlated expression between the pBS-Gln3 and pBS-Gcn4 reporters (Fig 4 E). Cells with high levels of pBS-Gcn4-tSc fluorescence also induce a high expression of pBS-Gln3-qV. This result disfavors a model where we have two subpopulations of cells: one that activates strongly Gcn4 and weakly Gln3 and another one where Gln3 is the main TF activated, while Gcn4 remains lowly active.

This correlated expression rather indicates that the whole cell population goes through a first phase of Gcn4 activity and subsequently activates Gln3. However, due to the long lifetime of FP proteins, it is difficult to know if Gcn4 and Gln3 can be active simultaneously in a cell or if there is a clear temporal separation between these two activities.

During the time-lapse, cells slowly deplete the ammonium available in the well at the onset of the experiment. In rich ammonium medium (200mM), this has little influence on the signaling activity of the cells because the N-source remains abundant throughout the experiment. In poor medium, the decrease in ammonium concentration in the environment triggers the switch between Gcn4- and Gln3- driven expressions. Based on our results, the transition occurs at a concentration between 50µM and 12.5 µM, because when the time-lapse starts with only 12.5µM NH_4_^+^, no induction of the pBS-Gcn4-qV is observed. Gcn4 and Gln3 are both regulated by TOR activity, yet via different mechanisms ^27,56^, which could explain why they are active in different concentration ranges. Interestingly, the temporal coupling between the Gln3 dependent expression and the change in eccentricity suggests that part of the morphogenetic change is controlled by proteins regulated by the TF Gln3 or by the same TOR-dependent molecular mechanisms that activate Gln3.

### Inhibition of the fgMAPK

While Gln3 and Gcn4 activations are evidently controlled by nutrient levels, the fgMAPK reporter based on Ste12/Tec1 is induced in rich and poor conditions. To verify the dependence of the pBS-Ste12/Tec1-qV reporter on the fgMAPK pathway activity during our microscopy assay, we mutated the MAPK Kss1 to generate an analog sensitive allele (*kss1-as*), where the kinase activity can be blocked by the addition of an ATP analog ^57,58^. Cells bearing the pBS-

Ste12/Tec1-qV reporter treated with the inhibitor 1-NMPP1 displayed a minimal increase in fluorescence when grown in SLAD, while the control sample treated only with DMSO showed the expected induction of the Ste12/Tec1 dependent reporter (Fig S7A). Using this strategy, we could verify that the fgMAPK contributed to the morphogenetic changes observed upon growth in SLAD. In cells treated with 1-NMPP1 the cellular eccentricity remained below 0.55 (Fig S7B).

In addition, we also tested if the signaling activity in the fgMAPK could modulate the activity of the other TFs implicated in FG. The inductions of the Gcn4-, Msn2- and Cat8-dependent reporters were significantly reduced in cells expressing the *kss1-as* and treated with 1-NMPP1 (Fig S8). Interestingly, inhibition of the fgMAPK had no significant influence on the TOR-dependent Gln3 reporter and on the second increase in eccentricity observed around the 6h time point. These results suggest that both Gln3 and the fgMAPK pathway contribute to the elongation of the cells in SLAD medium, with the fgMAPK controlling the early phase of elongation and Gln3 the later one (Fig S7B).

### fgMAPK activity in rich nutrient conditions

The essential contribution of the fgMAPK pathway to the filamentation response has been known for a long time ^35,59^. Because the FG phenotypes are primarily visible under nutrient limitation, it has been proposed that low nutrients stimulate the fgMAPK cascade ^59,60,9^.

However, in our experiments, we observe that the pBS-Ste12/Tec1 reporter is induced independently of nutrient levels, suggesting that alternative cues can activate the fgMPAK pathway.

The environment surrounding the cells may play a critical role in activating the fgMAPK pathway and we wondered whether we would observe different inductions between cells grown in a shaking culture compared to the surface of a well plate with liquid or agar medium. We performed confocal imaging of microcolonies after overnight growth in these different conditions (Fig 5 A and B). Cells grown in a well slide with SAD or SLAD in 2% agar or in liquid media were compared to cells grown in 5ml of SAD in a shaking culture tube. Low cell densities were used to inoculate the samples to ensure growth throughout the incubation time at 30° (∼16hrs). We cannot detect a statistical difference between samples incubated in wells with liquid or 2% agar with SAD or SLAD. However, cells grown in the shaking culture tubes expressed significantly lower levels of the pBS-Ste12/Tec1-qV construct than the cells grown in the wells, either in SAD liquid or in SLAD 2% agar.

**Figure 5.**
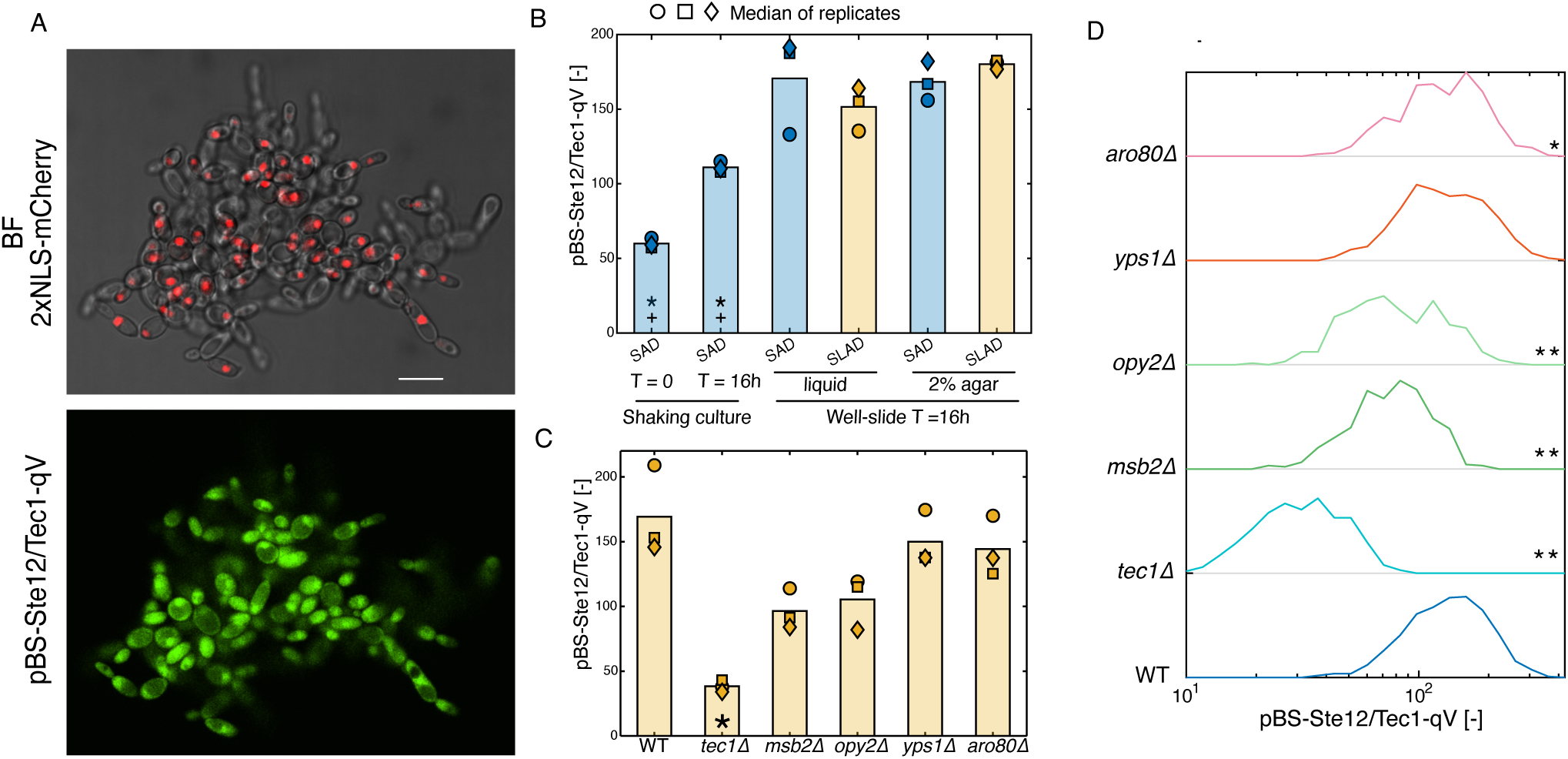
Induction of the fgMAPK pathway. A. Confocal image of a micro colony grown in SLAD 2% agar for 16 hours. The upper panel shows the transmission image and the red fluorescent nuclear tag. The lower panel displays the fluorescence level of the pBS-Ste12/Tec1-qV reporter. B. Average fluorescence of the pBS-Ste12/Tec1-qV reporter in WT cells grown in different conditions. The bar represents the mean of 3 biological replicates. The median of each replicate is represented by a marker. The stars and crosses indicate respectively statistical differences relative to the measurement performed in the well slide with SAD and liquid medium or with SLAD in 2% agar (t-test, p-value < 0.05). C. Average fluorescence of the pBS-Ste12/Tec1-qV reporter in WT and mutant cells grown in SLAD 2% agar for 18h. The bar represents the mean of 3 biological replicates. The median of the population for each replicate is represented by a marker. The star indicates a statistical difference relative to the WT response (t-test, p-value <0.05). D. Histogram of the average cellular fluorescence of the pBS-Ste12/Tec1-qV reporter in individual cells measured in WT and mutant cells after 18h of growth in SLAD 2% medium in a well slide. The star indicates that the histogram of the mutant is significantly different from the WT measurement (t-test, * p-value <0.05, ** p-value <10^-5^).

Two hypotheses could explain the higher level of induction of the pBS-Ste12/Tec1-qV reporter in wells compared to a shaking culture. The first one could be a local increase in the concentration of quorum sensing molecules in the vicinity of microcolonies. For instance, it has been suggested that small alcohols could stimulate FG induction ^61,62^. Alternatively, the shedding of mucin Msb2 could trigger FG signaling in neighboring cells ^60,63^. The second hypothesis is that contact with the surface of the well could induce a mechanical stimulus inside the cells, which could be relayed by the fgMAPK pathway.

Genetic perturbations in diploid cells were performed to test the influence of different genes on the pBS-Ste12/Tec1-qV expression output. As expected, removing the TF *TEC1* had a large impact on the expression of the reporter. However, the deletions of *ARO80* to block the production of aromatic alcohols and of *YPS1,* which is implicated in the cleavage of *MSB2,* result in minor changes in the expression of the reporter. These results suggest that quorum sensing is not triggering the activation of the fgMAPK pathway. On the contrary, deleting *OPY2* and *MSB2* lowered significantly the induction of pBS-Ste12/Tec1-qV reporter. Msb2, Opy2 and Sho1 are surface proteins that have been genetically mapped as the activators of the fgMAPK pathway ^9^. These proteins are also known to play a role in the activation of the HOG pathway in response to osmotic stress ^40,64^ and to confer resistance upon mechanical pressure ^65^. This suggests that the role of these well-known mechano-sensing proteins in the fgMAPK pathway could be to relay surface contact or mechanical strain stimuli.

### Pressure Sensing

To verify that the fgMAPK can be activated by mechanical cues, we used microfluidic confinement devices (Fig 6A) ^65^. Rich medium diffuses inside the chambers via small side channels, providing a constant supply of nutrients for cellular growth and division. The accumulated growth of the cells inside the main chamber induces a build-up of pressure (Fig 6A). Under these conditions, we detect a strong activation of the Ste12/Tec1-dependent reporter, while the control non-binding reporter is not activated (Fig6 B and C, Supplementary Movie 9, 10). The amount of pressure inside a chamber can be estimated by the level of deformation of the walls of the chamber induced by the cells. A linear correlation between the final pressure in the chamber and the level of pBS-Ste12/Tec1-qV induction is observed (Fig 6D). The physical forces exerted on the cells in the confinement chamber or in our well-slide experiments are radically different. However, this experiment confirms that the fgMAPK pathway can be activated by mechanical cues in a rich nutrient environment. Moreover, the tight correlation between the pressure in the chamber and fluorescence signal throughout the experiment demonstrates that the pathway is sensitive to a large pressure range (Fig S9), suggesting that it could also detect the much weaker stimuli generated by the surface contact.

**Figure 6.**
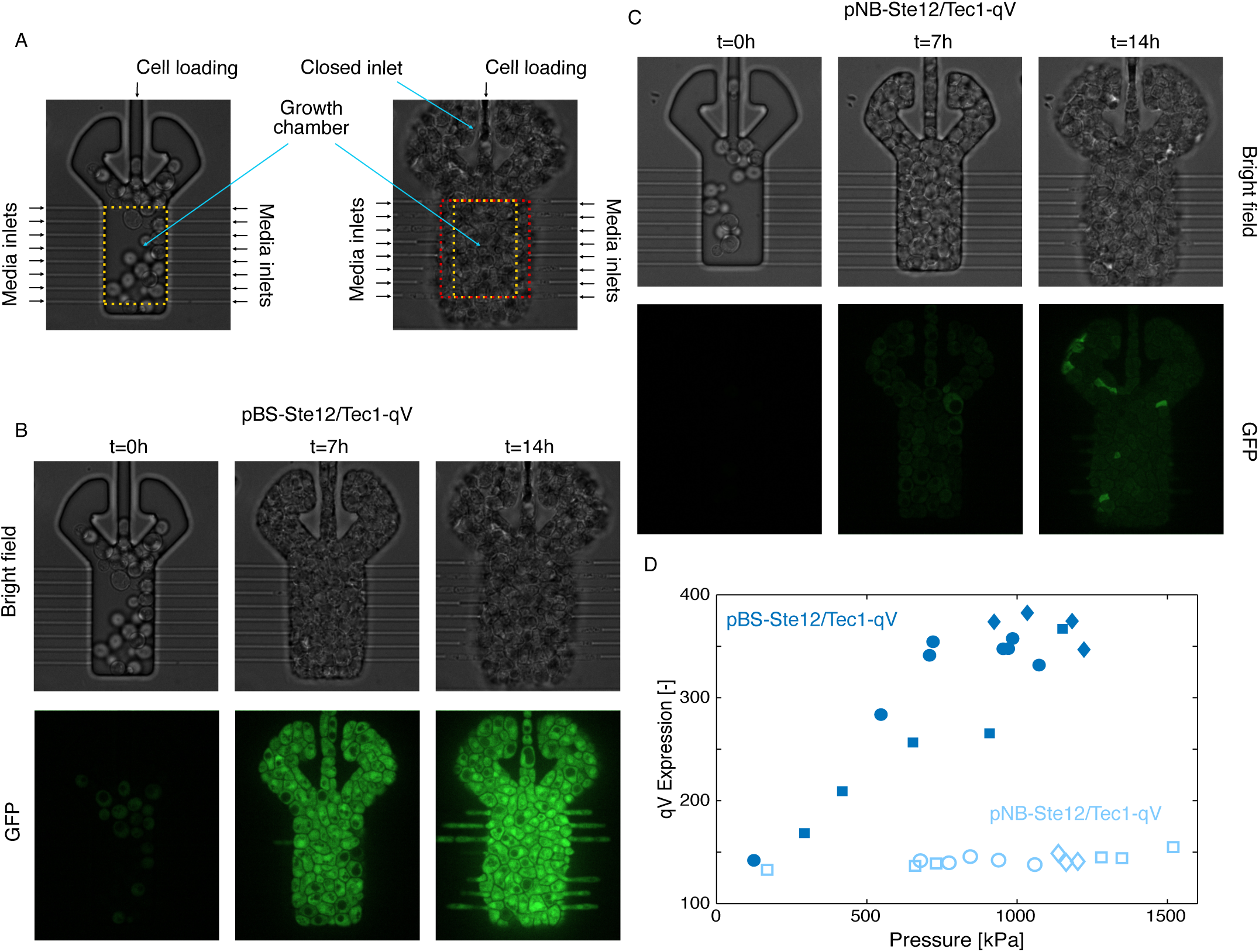
The fgMAPK pathway senses pressure. A. Image of the confinement chamber with the media and cell inlets highlighted by black arrows. Accumulation of cells in the growth chamber leads to a closing of the inlets (blue arrows) followed by a deformation of the PDMS substrate. The initial chamber size is depicted in yellow and the final size in red. The extent of the deformation can be linked to the level of pressure experienced by the cells in the device. B and C. Cells bearing the functional Ste12/Tec1 reporter (pBS-Ste12/Tec1-qV, A) and the mutated version (pNB-Ste12/Tec1-qV, B) were loaded into the confinement chamber and grown for 14 hours. The expression of the qV protein was monitored in the GFP channel. D. Correlation between the pressure accumulated in the chamber at the end of the experiment and the average fluorescence in the chamber at the same time (14 hours). The functional sensor is plotted in dark blue with closed markers, the mutated sensor in light blue with open markers. Each marker corresponds to the quantification of one confinement chamber, and the different marker shapes identify measurements obtained from different replicate experiments.

## Discussion

In this study, we used endogenous and synthetic promoters to monitor the dynamics of gene expression during the switch from vegetative to filamentous growth. While *FLO11* has been a hallmark of FG induced gene in numerous studies, in our experiments, its bimodal expression and the repression of expression at later time points don’t seem to provide a straightforward relationship between cellular morphology changes and gene expression induction. However, we identified p*YCT1* as an improved reporter of FG because it provides homogenous and sustained gene expression during the transition to FG. Interestingly, while at the population level, there is a correlation between the increase in fluorescence of the cells and their eccentricity, at the single cell level, we fail to observe such a relationship. This is also true for the pBS-Gln3-qV or the pBS-Ste12/Tec1-qV reporters. This is partly due to the uniform expression of the reporters within a microcolony. Cells in the center and at the periphery of a colony tend to express the reporter to a similar extent, while only the growing cells at the periphery alter their shapes and become elongated. In the future, it will be important to implement a tracking of the genealogy of the entire microcolony to infer the temporal link between the expression of the reporter and the modification of the cellular morphology in each cell.

One drawback of using endogenous reporters is their complex regulation by multiple TFs, making it difficult to isolate the contributions of the different signaling pathways implicated in the activation of the reporter. The engineering of synthetic promoters, which harbor binding sites for well-defined TFs allows to overcome this problem to some extent. This approach is ideally suited for Ste12/Tec1, which is specifically activated by the fgMAPK pathway. This strategy was equally successful for Gln3 because pBS-Gln3-qV was specifically activated in low ammonium conditions, implying a direct link to the repression of TOR activity.

Interpretation of the induction of synthetic promoters based on Msn2 or Gcn4 binding sites is more challenging because these TFs are known to respond to various cues and might thus depend on more than one signaling cascade. Msn2 is a well-identified target of the PKA pathway ^66^, but is also known to be activated by general stresses such as hyper-osmotic stresses, heat or rapamycin treatment ^23,67^. Gcn4 is activated by amino-acid starvation and can be controlled by the TOR pathway in low nitrogen conditions and in low glucose via the action of PKA. Moreover, it has been shown that there is extensive cross-talk between nutrient signaling pathways and notably between TOR and PKA ^69^. Therefore, extrapolation of a synthetic expression reporter signal to an upstream activating signaling cascade has to be performed cautiously.

We have shown the fgMAPK pathway is also enhancing the nutrient signaling response in SLAD. Blocking Kss1 activity dampens the response from the Gcn4, Cat8 and Msn2 -dependent reporters. However, it remains to be determined at which level these cross-activations occur. It could be at the signaling level where Kss1 directly modulates the activation of members of nutrient signaling pathways. Alternatively, Kss1, via Ste12/Tec1-dependent gene expression, could modulate the level of the proteins implicated in these pathways. In any case, these results reinforce the idea that individual signaling cascades do not act in isolation in the cell but form a complex signaling network. Identifying the molecular mechanisms of these cross-inhibition and cross-activation remains challenging. Synthetic expression reporters can contribute to solving this riddle. However, signaling activity reporters that provide direct readout of kinase activity may be better suited for this task ^70^.

While numerous studies of filamentous growth in *S. cerevisiae* have focused on endpoint measurements after multiple days in nutrient limiting conditions, we have developed assays to monitor the first 10 hours of FG induction under the microscope. To simplify the interpretation of the results, we have focused our study on the limitation of glucose, ammonium or both. In all three cases, we observe some level of FG induction, although the morphology of the cell and the microcolony they form are slightly different. Under the experimental conditions tested, we have observed low levels of induction from the SNF and PKA pathways using the Msn2, Flo8 or Cat8 dependent reporters. Both SNF and PKA have a well-recognized role in FG induction ^13,71^. We hypothesize that these signaling pathways could play a more important role in a later phase of the process, when glucose has reached very low levels and alternative C-sources in the medium start to be utilized by the cells.

While previous studies have suggested that low nutrients were triggering the fgMAPK cascade ^14,34,35^, in this study, we demonstrate that the fgMAPK is induced independently of nutrient levels. We rather propose that the fgMAPK senses mechanical signals and is activated when cells grow on a solid surface. The previously identified role in force sensing of the surface proteins Msb2, Opy2 or Sho1 at the helm of the fgMAPK pathway aligns well with this hypothesis. Based on our results, we envision the following model for the induction of FG in SLAD medium. In cells in liquid and nutrient rich environments, the fgMAPK is not stimulated, while TOR activity blocks Gln3 and Gcn4-dependent expression. If the cell falls on a surface, the fgMAPK pathway will be stimulated. Subsequently, consumption of the available pool of nitrogen lowers the TOR activity, which enables the activation of Gcn4. Further depletion of the N-source limits even more the TOR activity and Gln3 starts to be active. The combined activity of the fgMAPK and of Gln3 triggers the induction of filamentous growth (Fig 7).

**Figure 7.**
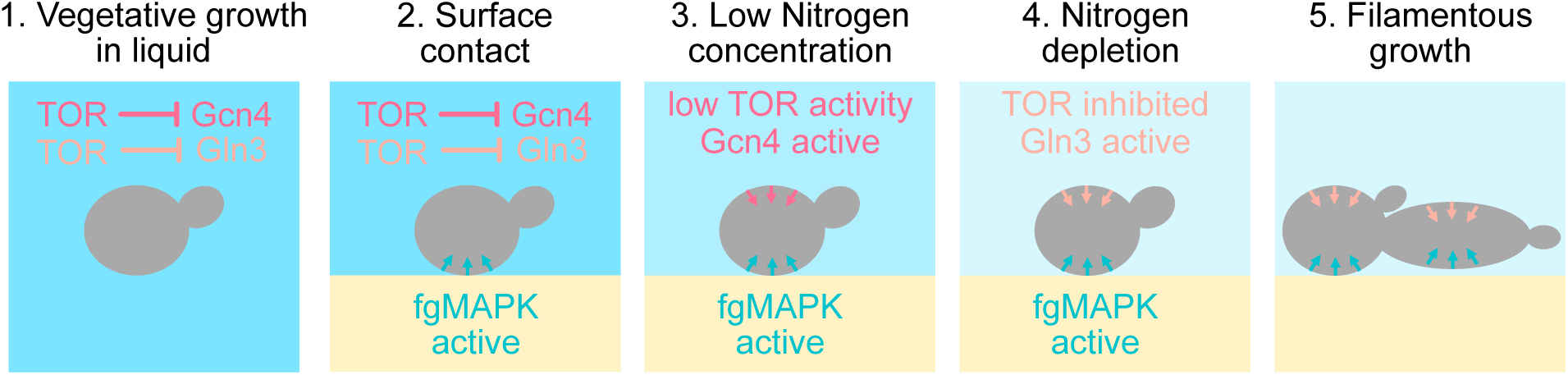
Model of filamentous growth induction by low ammonium. The different steps towards FG in SLAD are depicted. 1. Planktonic cells grown in a rich liquid medium have a full active TOR pathway that represses Gln3 and Gcn4. 2. As cells land on a solid substrate, the contact with the surface activates the fgMAPK pathway. 3. In parallel, the low level of ammonium results in the partial inhibition of TOR, which allows Gcn4 to become active. 4. Subsequent depletion of the nitrogen source further limits TOR activity and allows Gln3 to initiate its transcriptional program. 5. Combined fgMAPK and Gln3 activities promote the entry into the filamentous growth regime, characterized by elongated cells and a polarized division pattern.

Despite their thick cell walls, fungi have developed sophisticated means to transduce mechanical cues ^72^. For instance, fungal plant pathogens have devised numerous strategies to detect the surface of leaves, and part of this signal transduction is performed by MAPK pathways connected to the surface sensors Sho1 and Msb2 ^73^. In the human pathogen, *Candida albicans*, detection of surface contact is transduced via the MAPK Mkc1 and contributes to hyphal growth and biofilm formation ^74^. The membrane sensors Sho1, Msb2 and Opy2 have homologues in *C. albicans,* where they regulate the activation of Cek1, a MAPK which is implicated in invasion and biofilm formation. Mkc1 and Cek1 are respectively the homologues of Slt2 involved in the cell wall integrity pathway and Fus3 controlling the mating response in *S. cerevisiae*. Neither Fus3 nor Slt2 contribute to FG, suggesting that the surface sensing that is required for FG activation in many fungal species might be transduced by Kss1 and the fgMAPK pathway in *S. cerevisiae*.

## Materials and methods

### Yeast strain and plasmids

The plasmids and strains used in this study are listed in the Supplementary File S1. The synthetic reporters were cloned in a pSIV-URA vector ^53^. The quadruple Venus (qV) is cloned between HindIII and XhoI and the *SIF2* terminator (427 bp) is cloned between XhoI and KpnI. The synthetic promoter consists of the core promoter of *CYC1* (183 bp) and an upstream activator region containing the binding sites. Supplementary File S1 contains a list of the characteristics and sequences of the tested promoters. The endogenous *FLO11* (-3200 to 0) and *YCT1* (-1000 to 0) promoters were cloned between SacI and HindIII.

The plasmids with the synthetic reporter systems were transformed in diploid strains of the Σ1278b background with *URA3* deficiency ^72^. A nuclear marker was also added to the strain by integrating a 2A-2xNLS-mCherry:NAT cassette downstream of the *LYS21* gene.

The diploid gene deletions were obtained by performing deletions in haploid cells sporulated from the parental strain containing the pBS-Ste12/Tec1-qV reporter and where the nuclear marker cassette was switched from NAT to KAN. The entire locus of the gene of interest was deleted by a NAT cassette ^73^. A MATα strain with the pBS-Ste12/Tec1-qV, the nuclear marker, and the deleted gene was mated to a MATa cell with the gene deletion to obtain the diploid homozygote deletion mutant.

The *kss1-as* strains were generated by CRISPR-Cas9 engineering using a modified version of the Cas9 sgRNA plasmid ^77^. The modified plasmid contains a NAT resistance marker, the Cas9 is expressed from a strong constitutive promoter (p*TDH3*) and the guide RNA is expressed from the p*SNR52* promoter. The original cloning strategy has been modified to allow the digestion of the backbone with AatII and XbaI for the insertion of the selected guide RNA. Two 120 bp oligos were annealed and amplified with polymerase to generate a 225bp double stranded repair DNA fragment with 60 bp homology to the *KSS1* sequence on both ends and while the central part contains the E93A mutation and a codon shuffled sequence to prevent annealing the guide RNA once the repair has occurred (Supplementary File 1). After transformation, the colonies were genotyped by sequencing a 500 bp region containing the mutated sequence.

The endogenous tagging of *FLO11* was also performed by CRISPR-Cas9 engineering. The mCitrine and mCherry tags were introduced in a haploid MATα and MATa strain with Hta2- tdiRFP::KAN markers. The fluorescent proteins were inserted between P222 and V223, after the A domain and before the first repeat of the B domain. The *FLO11* locus was targeted with a guide RNA at position 623 (P208). The guide RNA was cloned in a Cas9 containing plasmid similar to the one described above with a URA3 auxotrophy selection marker. The repair DNA was generated by amplifying the selected fluorescent protein by PCR using primers that contained 100bp overhangs. The forward primer contained a homology region of 55bp and a 45 bp sequence with a codon shuffled sequence replacing the original sequence between P208 and P222 (Supplementary File 1). The transformants were screened by microscopy for a fluorescence signal at the cell periphery, then genotyped by PCR to verify the size of the repeat sequence region and the PCR was sequenced to verify the correct insertion of the FP. The positive transformants were also tested for invasion. Note that tagging of the FLO11 locus results in a hypomorphic allele with reduced invasion capabilities. After obtaining the MATα tagged strain, the antibiotic marker cassette was replaced by a *URA3* auxotrophy cassette. The MATa and MATα strains were mated and the diploid was selected on SD-URA and YPD+KAN plates.

The dual reporter strain for Gln3 and Gcn4 activity was built in a Σ1278b strain with *HIS3*, *URA3* and *LEU2* auxotrophy ^75^. The nucleus is marked with a Hta2-CFP:*HIS3* tag. The pBS- Gln3-qV is integrated in the *URA3* locus and the pBS-Gcn4-tSc (triple-Scarlet-I3^79^) is inserted in the *LEU2* locus.

To assess that the filamentous characteristics of the transformed strains were not altered, the invasion capacity of the strain was assessed. A set of 8 colonies was grown in 3ml YPD and 5µl of the overnight culture was spotted on a YPD plate and incubated for 5 days at 30°. The colonies were washed away to monitor the invasion scar generated by each transformant. The parental strain was spotted in the middle of the plate for comparison. For WT strains, the transformant similar to the WT control was selected. For mutant strains, comparisons between the transformants were performed and strains with an outlier phenotype were discarded.

### Time-Lapse Imaging

The strains were grown overnight in SAD medium (low fluorescence Yeast nitrogen Base without ammonium sulfate, YB (ForMedium, CYN6510), 2% glucose 20 mM ammonium sulfate), diluted in the morning to OD 0.025, grown the entire day and diluted in the evening to ensure log-phase growth until the next morning. Unless specified in the figure legend, this 24- hour log-phase growth was required to dilute fluorescent proteins that can be expressed during the first overnight growth to saturation (for instance, pBS-Msn2-qV or pBS-Cat8-qV in Figure S4). The OD of the 5 ml log phase culture was measured. The cells were filtered on a cellulose filter (Millipore, HAWP 02500) washed twice with 5ml YB (ForMedium, CYN6510). The filter with the washed cells was collected from the filtering apparatus and placed in 5ml YB. The resuspended cells were diluted to OD 0.01 in 1 ml YB according to the OD measured before the filtration. Fifty microliters of the diluted washed cells were deposited at the center of the well of a 96-well plate (SwissCI, PS96B-G175) previously coated with ConcanavalinA (Sigma-Aldrich, C2010). Cells were left to settle for 20min in the incubation chamber of the microscope set at 30°. Then, 400µl of molten agar medium was deposited drop by drop on top of the cells. Low-melting agar medium (Eurobio, GEPAGA04-62) was prepared before the filtration of the cells by dissolving 0.1 g of low-melting agar in 10ml YB by heating slowly in the microwave. One milliliter aliquots were prepared. In each microtube, the desired glucose and ammonium concentration were defined by adding the proper volume of 50% glucose and ammonium sulfate at 0.5M or 20mM to reach final concentrations of 2% (D) or 0.05% (LD) and 20 mM (A) or 50 µM (LA). The aliquots were kept at 40° in a heat block before being loaded into the well. After loading the samples, the wells were covered with parafilm and the lid of the 96-well-plate to limit the desiccation of the agar medium.

Cells were imaged on an epifluorescence microscope (Nikon, Ti2-eclipse) controlled by micro-manager ^77^ using a 40x air objective. A Lumencor Spectra III light source was used to excite the sample using a multi-band dichroic filter and emission filters to detect the YFP and RFP signals with a sCMOS camera (Hamamatsu Fusion BT). In parallel, a single bright field transmission image was recorded. Cells were imaged every 10 or 15 min for up to 12 hours. Typically, 5 positions per well were recorded and up to 24 wells with different media conditions or strain backgrounds were imaged in parallel.

### Confocal Imaging

A similar protocol was used to prepare the samples for the end-point confocal imaging measurements. Cells were grown overnight to saturation in SAD medium and grown for 4 to 6 hours in 5 ml SAD to reach an OD around 0.1. Cells were filtered, as described above, and resuspended in YB and diluted to an OD of 0.001 in 1 ml. This low cell density allows cells to have enough nutrients to sustain growth throughout the overnight incubation and into the next morning when they were imaged. Fifty microliters of the diluted cells are loaded into a well.

After 20 min of settling time on the bench, the 400 µl of the desired medium kept at 40°C (low-melting agar or liquid) was slowly added to the desired well. The wells were covered with parafilm and the lid of the 96-well-plate and placed in a 30°C incubator.

After 16 to 18 hours of incubation, the cells were imaged on a confocal microscope (Nikon, Ti2 AxR) using a 40x air objective using the 514nm laser at 20% power. The YFP signal from the qV expression reporter was collected in a 524-551 nm range, while the RFP signal of the nuclear tag was measured between 562 and 634 nm. The 2048x2048 pixel image was scanned at 0.8ms/pixel with four times averaging with a 0.109 µm pixel size. The pinhole was open to 0.5 Airy unit, defining a Z-slice of 1.06 µm at 514 nm to strictly limit the contribution of out-of-focus cells to the fluorescence signal. Ten fields of view per condition were recorded corresponding to one (sometimes two) micro colony.

### Compression Chamber

The experiments have been performed according to the previously described protocol ^65^. The cells are loaded into the chambers and imaged for 14 hours every 5 minutes in the bright field and GFP channels. SD-full medium is fed from the lateral media channels to allow constant cellular growth. As the cells fill the chamber, they will accumulate inside the chamber, close the inlet and then build-up pressure. The average fluorescence signal from the chamber was quantified at each time point in the GFP channel. The width of the chamber is measured from the bright field image. To convert the chamber width into a pressure value, a calibration was performed where the chambers were inflated with pressurized air and the deformation of the chambers was measured.

### Image Analysis

Images were analyzed using an update version of the YeastQuant platform ^81^. The cell contours were segmented by CellPose 2.0 ^82^ using a combination of the bright field and nuclear fluorescence images. The neural network was trained on images of filamenting cells to improve the detection of elongated cells, which were poorly segmented with the original cyto2 pipeline. The segmented cells contours are subsequently fed to the YeastQuant pipeline and combined with a segmentation of the fluorescent image of the nuclei to define the cells of interest and eliminate out of focus cells or other objects recognized by CellPose. Fluorescence intensity and shape measurements are performed on the segmented cells. Due to the difficulty in tracking the fate of individual cells in the microcolonies, we plot for each time point the mean, median or percentiles of all the cells identified by YeastQuant. At the onset of the time-lapse, this can be as few as 50 cells for one condition, but it will reach 500 to 1000 after 8 hours of growth.

A similar strategy was used to segment cells in the microcolonies imaged with the confocal microscope, however, only the CellPose mask obtained with the cyto2 model was used to define the segmented cells. Subsequently, we used the signal from the nuclear tag in the RFP channel to select cells which had the nucleus in the focal plane. Combining the mean intensity of the 20 brightest pixels in the cell object and the normalized standard deviation of the RFP signal in the cell object allowed to reject out of focus cells.

Dedicated scripts were written in Matlab (R2024b, Mathworks) to analyze and plot the data obtained from the YeastQuant pipeline.

## Acknowledgments

Research in the Pelet lab is funded by the Swiss National Science Foundation (Grant N°: 320030-231544) and the University of Lausanne. We thank all members of the Pelet Lab for helpful discussions. Nienke Jager, Stella Parzanese, Vanissa Froissard and Thibo Bellanger provided technical help to generate strains and plasmids for the study. We want to thank Matthias Peter at the ETH in Zürich and Claudio De Virgilio at the University of Fribourg for sharing yeast strains and Aleksandar Vještica for helpful discussions.

HK and MD would like to acknowledge the ERC Starting Grant UnderPressure (Grant agreement N°: 101039998). Views and opinions expressed are, however, those of the author only and do not necessarily reflect those of the European Union or the European Research Council.

Neither the European Union nor the granting authority can be held responsible for them.

## Author Contributions

SPa and SPe designed the study and performed the epifluorescence time-lapse measurements. Spa and DS established the live-cell FG assay. ML performed the confocal experiments. HK and SPa performed the microfluidic confinement experiments. DS generated the synthetic promoter plasmids and strains. VB constructed the *FLO11* internal tagging strains. YD and VV generated strains and plasmids for the study. SPe and MD acquired funding. SPa and SPe analyzed the data and wrote the original version of the manuscript. All authors contributed to the final version of the article.

