## Supplementary Figures for "Cellular integration of surface and nutrient signals during the early stage of filamentous growth in *Saccharomyces cerevisiae*"

### Supplementary Figures Legends

*Supplementary Figure 1. Signaling pathways implicated in filamentous growth.*

Scheme of the main signaling pathways implicated in the induction of filamentous growth. Sensing of the carbon source is performed by the PKA and SNF pathways, while the Nitrogen-source is sensed by TOR. Low amino-acid levels will trigger the activity of Gln2 and Gcn4. The fgMAPK plays an important role if FG induction. However, it remains unclear how nutrient signals are sensed by the membrane sensors Msb2, Sho1 and Opy2 which have been shown to promote the activity of the fgMAPK pathway.

*Supplementary Figure 2. Experimental process*

A. Scheme of the preparation of cells for the FG induction experiments. Cells are grown in log-phase prior to filtration and re-suspension in a medium without nutrients (YB).

B. Low-melting agar is prepared in YB, aliquoted and the desired volumes of glucose and ammonium sulfate are added to the microtube to define the amount of nutrients in the medium. The low-melting agar is then kept at 40°C before use.

C. For epi-fluorescence time-lapse imaging, the filtered cells are diluted to OD 0.01 and loaded into the well of a 96-well plate. After 20min of settling time, 400µl of molten agar is added to the well. The well is closed with parafilm and the lid of the well plate. The time-lapse imaging can start to follow the growth of the cells for up to 12 hours.

D. For confocal end-point imaging, the filtered cells are diluted to OD 0.001 and loaded into the well of a 96-well-plate. After 20min of settling time, 400µl of molten agar is added to the well. The well is closed with parafilm and the lid of the well plate. The well plate is incubated at 30°C overnight and imaged the next day after 16 to 18 hours of growth.

*Supplementary Figure 3. Endogenous FLO11 fluorescent reporter*

A. Structure of the FLO11 locus. The FLO11 has three domains A (for adhesion to surfaces and self-interaction) B (formed by tandem repeats) and C (unknown function). The sequence starts with a signaling peptide (S) for proper export of the protein and ends with a GPI anchor motif (G). Due to these last two features, N- or C-terminal tagging results in a non-functional protein. The transcription of the FLO11 locus is regulated by two long non-coding RNAs (ICR1 and PWR1), which control the bistable expression of the locus. Fluorescent proteins were inserted by CRISPR-Cas9 engineering between the A and B domains.

B. Images of the expression of the Flo11-mCherry and Flo11-mCitrine in their native loci. The scale bar represents 5 µm.

C and D. Dynamics of fluorescence apparition at the periphery of the cells in the yellow (C) or red (D) fluorescent channel. The dots represent the average cellular intensity from individual cells and the solid line represents the mean of the population. The percentage of expressing cells in the population is indicated. The threshold of expression is depicted by the dashed line. For clarity, up to 200 individual cells are displayed.

F. Correlation between the Flo11-mCherry and Flo11-mCitrine expression in individual cells at 2 4 and 8 hours after the start of the imaging. The cells which express both reporters and in orange, while the ones expressing only the Flo11-mCitrine or the Flo11-mCherry are in yellow and red, respectively. Non-expressing cells are in gray. The thresholds for expression are visualized by the dashed lines.

*Supplementary Figure 4. Comparison of the binding and non-binding reporters in SLAD medium.*

A , B , C and D. Dynamics of fluorescence apparition from the binding (yellow) and non-binding (gray) Gcn4 (A), Msn2 (B) Flo8 (C) and Cat8 (D) -dependent reporters in cells grown in SLAD agar. For Cat8, two promoters were constructed. pBS-Cat8$_{weak}$-qV (red) is based on the endogenous 72 bp sequence of the MLS1 promoter, while pBS-Cat8-qV contains the same MLS1 sequence with two strong consensus sites (TCCATTCATCCGA) displaying a stronger activation pattern. The dashed lines represent the median of the population and the shaded areas, the 10 and 90 percentiles. Note that for Panels B, C and D, the high fluorescence at the beginning of the time course is due to residual fluorescence from the overnight growth to saturation, which precedes the 4 hours of growth in SD-full before the filtration of the cells for the preparation of the samples for the time-lapse experiment. Due to this experimental artefact, the differential expression levels calculated in Figure 2C correspond to the difference between the maximum and the minimum of each curve.

*Supplementary Figure 5. Testing of expression reporters in various environmental stresses.*

A, B and C. Influence of Rapamycin (3 µM) on the induction of the Gln3 (A), Gcn4 (B) and Ste12/Tec1 expression reporters 4 hours after induction. Both Gln3 and Gcn4-dependent reporters display a clear induction of fluorescence when treated with rapamycin, demonstrated their dependence on TOR activity.

D, E, F and G. Induction of the Ste12/Tec1 (D), Msn2 (E), Flo8 (F) and Cat8 (G) expression reporters in SAD-Agar, 0.4 M NaCl, 2% Glycerol or in stationary phase. pBS-Msn2-qV displays the expected induction in osmotic stress medium (0.4M NaCl) and in stationary phase. pBS-Cat8-qV is induced in glycerol containing medium. pBS-Flo8-qV is slightly induced in stationary phase cells.

In all the above graphs, the square represents the mean of the population and the dots the individual cell measurements.

*Supplementary Figure 6. Characterization of reporters in four different nutrient environments.*

A. Evolution of the cellular area for cells grown in four different media. The solid line represents the mean of 9 curves from the three biological replicates performed with three different strains. The shaded area represents the standard error of the mean.

B, C and D. Dynamics of fluorescence levels for cells bearing the Msn2 (B), Cat8 (C) and Flo8 (D) -dependent expression reporters in the four media conditions. The dashed line and the markers indicate the mean of the population of cells measured at each time point.

E, F and G. Correlation between the eccentricity of the cells and the average cellular fluorescence of the pBS-Gln3-qV (E), pBS-Gcn4-qV (F) and pBS-Ste12/Tec1-qV (G). The triangles represent the average eccentricity and intensity of the population at specific time points. The dots represent the value of these two measurements in individual cells at 0h (black) and 9h (yellow). For clarity, up to 200 individual cells are displayed.

*Supplementary Figure 7. Inhibition of MAPK activity.*

A. Cells expressing an analog sensitive allele of the MAPK Kss1 are treated with the inhibitor NMPP1 (dashed, dark blue) or the vehicle DMSO (solid, light blue). The dynamics of the fluorescence intensity generated by the qV protein under the control of the synthetic Ste12/Tec1 reporter in cells grow in SLAD agarose. The median of the population and the 10 to 90 percentiles are indicated by the dashed line and the shaded area, respectively.

B. Influence of Kss1 inhibition on the evolution of the cellular eccentricity of cells grown in SLAD with the inhibitor NMPP1 (dashed, dark blue) or the vehicle DMSO (solid, light blue). The solid line represents the mean of 9 curves from the three biological replicates performed with three different strains. The shaded area represents the standard error of the mean.

*Supplementary Figure 8. Effect of Kss1 inhibition on other signaling pathways during FG induction.*

A. Differential expression of the qV construct promoted by synthetic promoters responding to the defined TFs in strains expressing the *kss1-as* allele. The bar represents the mean of 3 to 4 biological replicates (indicated by the markers). The star indicates that the measurements in the NMPP1 treated samples are significantly different than DMSO control (t-test, p-value <0.05).

B, C, D, E and F. Dynamic response of the pBS-TF-qV reporters in the presence of NMPP1 (dashed line) or DMSO (solid line), as a control, in the strains bearing the *kss1-as* allele. Cells are grown in SLAD-agar and the inhibitor (50µM) or DMSO (9% in YB) is added on top of the agarose at the beginning of the time-lapse measurements to reach a final concentration of 5µM. The dashed line and the marker represent the median of the population and the shaded area the 10 to 90 percentiles**.** Cells were cultivated for 8 hours in liquid SD-full medium before the preparation of the sample for the microscopy.

*Supplementary Figure 9. Temporal evolution of fluorescence and pressure during the confinement of the cells.*

A. Level of GFP intensity measured from the pBS-Ste12/Tec1-qV reporter in five different confinement chambers. The average fluorescence of the area where the cells are growing is plotted.

B. Evolution of the pressure extrapolated from the chamber width as a function of time. The colors represent the same chambers plotted in panel A.

C. Correlation between the pressure and mean chamber intensity during the 14 hours of the confinement experiment.

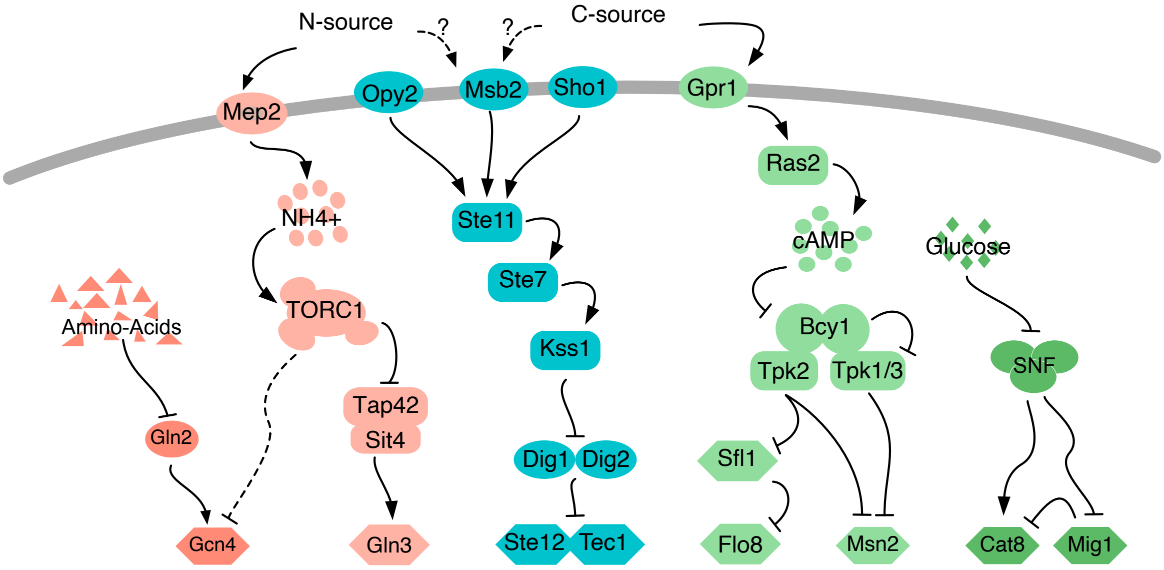
N-source
?
?
C-source
Mep2
Opy2
Msb2
Sho1
Gpr1
NH4+
Ste11
Ras2
Ste7
cAMP
Glucose
Amino-Acids
TORC1
Kss1
Bcy1
Tpk2
Tpk1/3
SNF
Tap42
Sit4
Gln2
Dig1
Dig2
Sfl1
Gcn4
Gln3
Ste12
Tec1
Flo8
Msn2
Cat8
Mig1

Figure S1

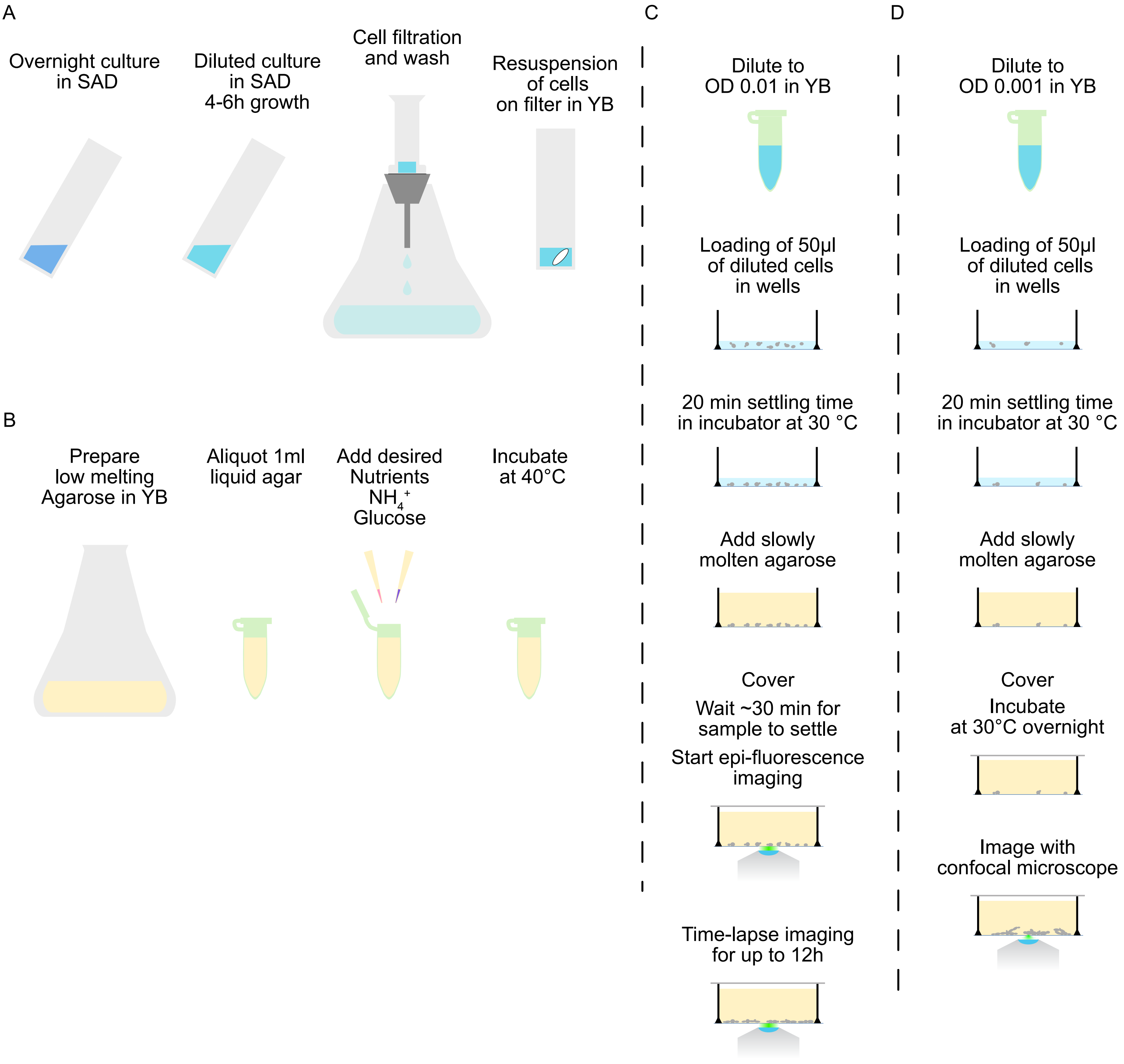
A
Overnight culture
in SAD
Diluted culture
in SAD
4-6h growth
Cell filtration
and wash
Resuspension
of cells
on filter in YB
B
Prepare
low melting
Agarose in YB
Aliquot 1ml
liquid agar
Add desired
Nutrients
$NH_4^+$
Glucose
Incubate
at 40°C
C
Dilute to
OD 0.01 in YB
Loading of 50µl
of diluted cells
in wells
20 min settling time
in incubator at 30 °C
Add slowly
molten agarose
Cover
Wait ~30 min for
sample to settle
Start epi-fluorescence
imaging
Time-lapse imaging
for up to 12h
D
Dilute to
OD 0.001 in YB
Loading of 50µl
of diluted cells
in wells
20 min settling time
in incubator at 30 °C
Add slowly
molten agarose
Cover
Incubate
at 30°C overnight
Image with
confocal microscope

Figure S2

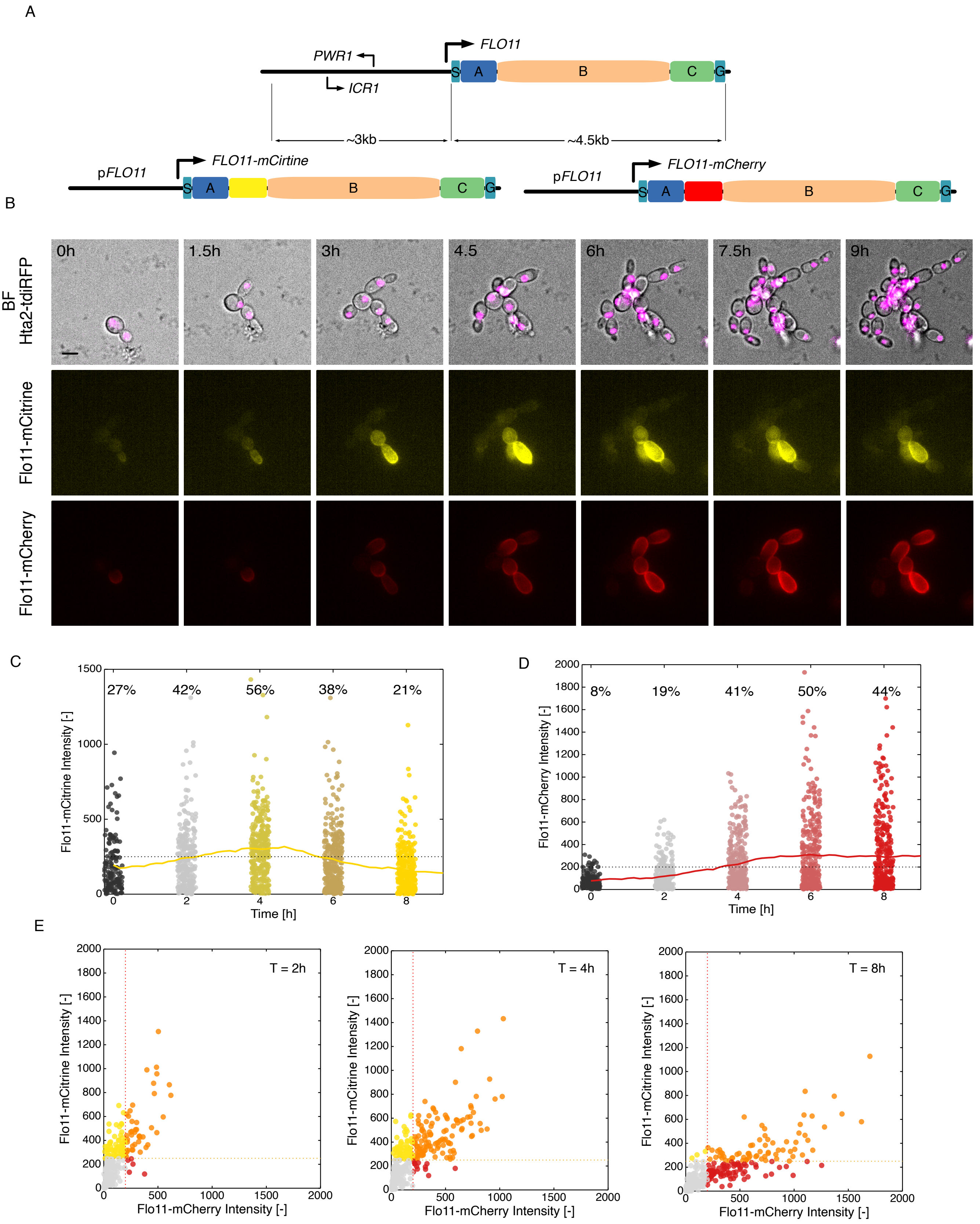
A
PWR1
ICR1
FLO11
S
A
B
C
G
~3kb
~4.5kb
pFLO11
FLO11-mCirtine
FLO11-mCherry
B
BF
Hta2-tdiRFP
Flo11-mCitrine
Flo11-mCherry
0h
1.5h
3h
4.5
6h
7.5h
9h
C
27%
42%
56%
38%
21%
Flo11-mCitrine Intensity [-]
Time [h]
D
8%
19%
41%
50%
44%
Flo11-mCherry Intensity [-]
Time [h]
E
T = 2h
T = 4h
T = 8h
Flo11-mCitrine Intensity [-]
Flo11-mCherry Intensity [-]

Figure S3

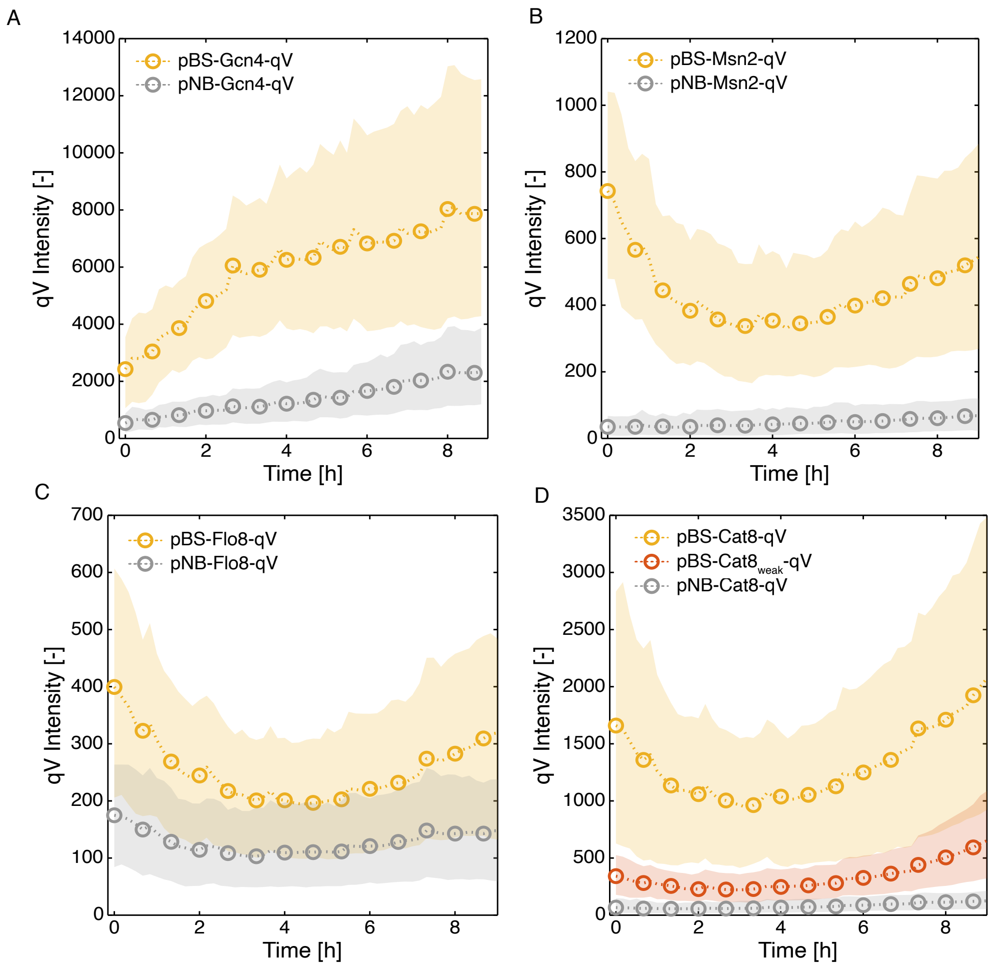
A
pBS-Gcn4-qV
pNB-Gcn4-qV
qV Intensity [-]
Time [h]
B
pBS-Msn2-qV
pNB-Msn2-qV
qV Intensity [-]
Time [h]
C
pBS-Flo8-qV
pNB-Flo8-qV
qV Intensity [-]
Time [h]
D
pBS-Cat8-qV
pBS-Cat8weak-qV
pNB-Cat8-qV
qV Intensity [-]
Time [h]

Figure S4

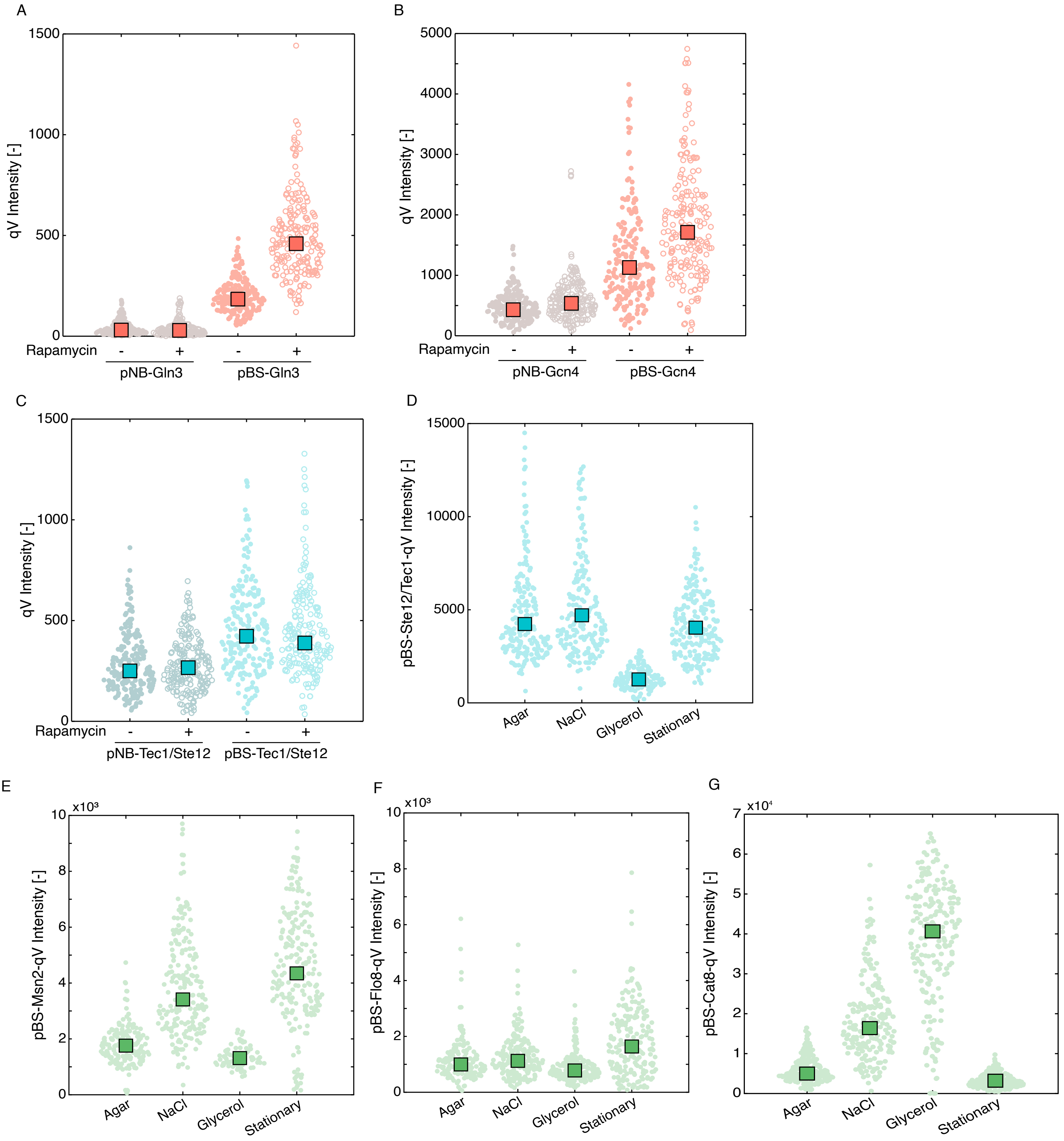

A
1500
1000
500
0
qV Intensity [-]
Rapamycin
- + - +
pNB-Gln3
pBS-Gln3
B
5000
4000
3000
2000
1000
0
qV Intensity [-]
Rapamycin
- + - +
pNB-Gcn4
pBS-Gcn4
C
1500
1000
500
0
qV Intensity [-]
Rapamycin
- + - +
pNB-Tec1/Ste12
pBS-Tec1/Ste12
D
15000
10000
5000
0
pBS-Ste12/Tec1-qV Intensity [-]
Agar
NaCl
Glycerol
Stationary
E
x10³
10
8
6
4
2
0
pBS-Msn2-qV Intensity [-]
Agar
NaCl
Glycerol
Stationary
F
x10³
10
8
6
4
2
0
pBS-Flo8-qV Intensity [-]
Agar
NaCl
Glycerol
Stationary
G
x10⁴
7
6
5
4
3
2
1
0
pBS-Cat8-qV Intensity [-]
Agar
NaCl
Glycerol
Stationary


Figure S5

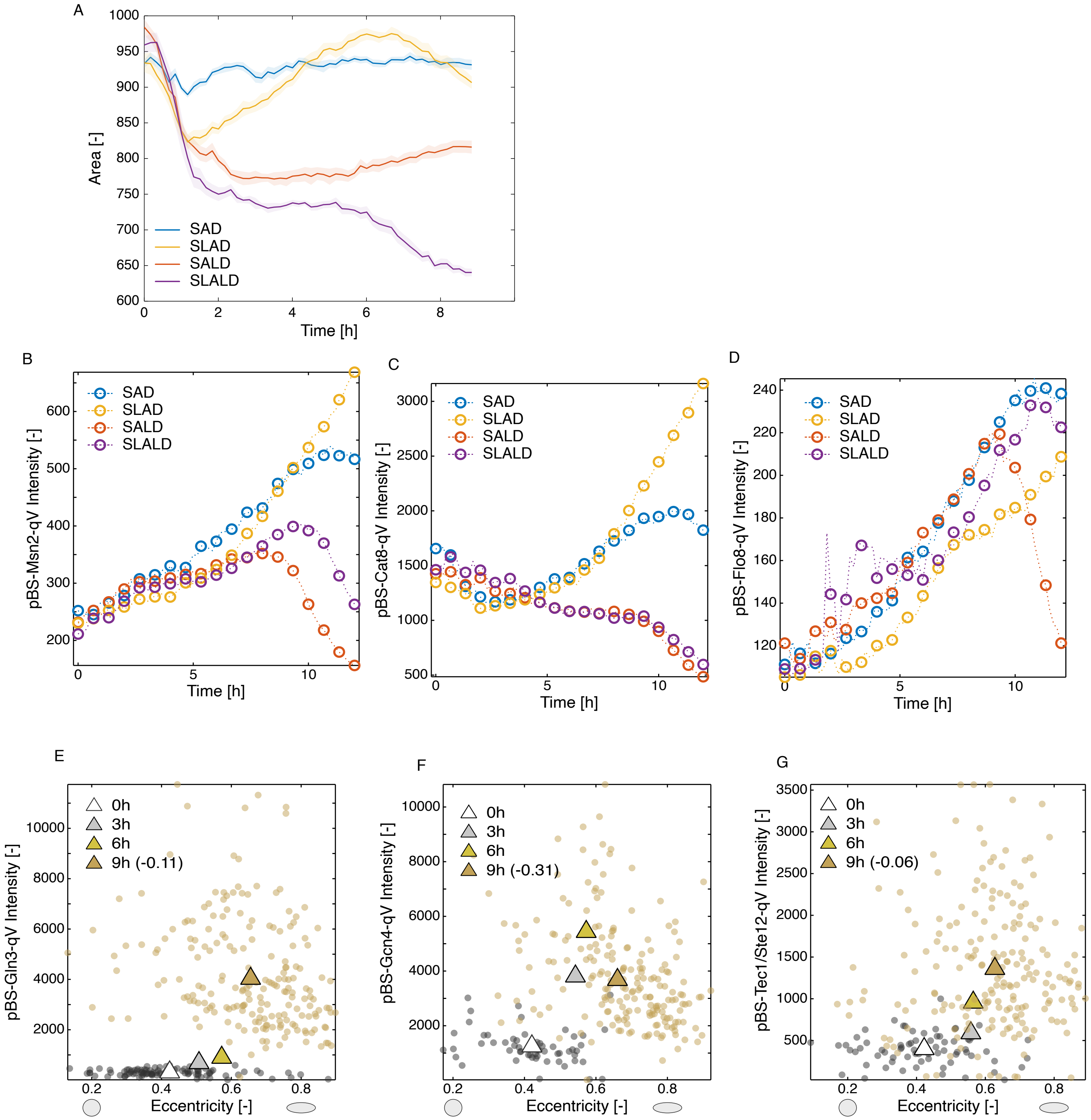
A
Area [-]
Time [h]
SAD
SLAD
SALD
SLALD
B
pBS-Msn2-qV Intensity [-]
Time [h]
C
pBS-Cat8-qV Intensity [-]
D
pBS-Flo8-qV Intensity [-]
E
pBS-Gln3-qV Intensity [-]
0h
3h
6h
9h (-0.11)
Eccentricity [-]
F
pBS-Gcn4-qV Intensity [-]
9h (-0.31)
G
pBS-Tec1/Ste12-qV Intensity [-]
9h (-0.06)

Figure S6

A

B

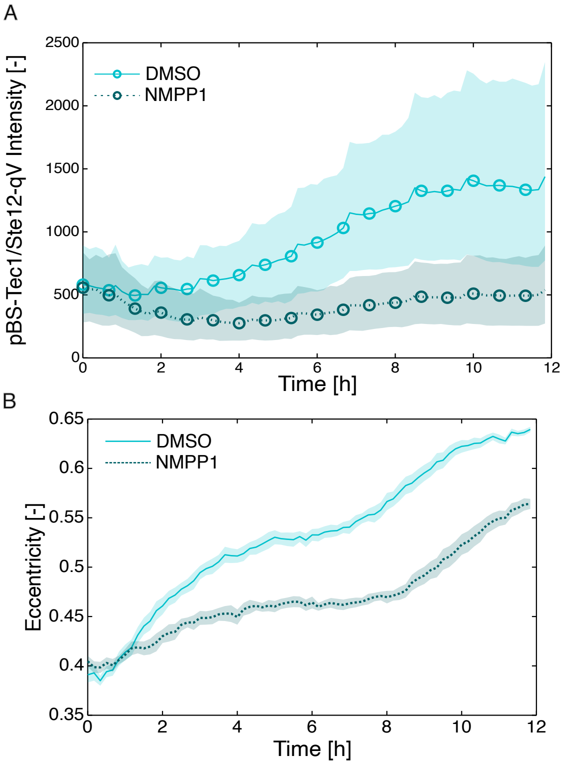

2500
2000
1500
1000
500
0
pBS-Tec1/Ste12-qV Intensity [-]
DMSO
NMPP1
0
2
4
6
8
10
12
Time [h]
0.65
0.6
0.55
0.5
0.45
0.4
0.35
Eccentricity [-]
DMSO
NMPP1
0
2
4
6
8
10
12
Time [h]


Figure S7

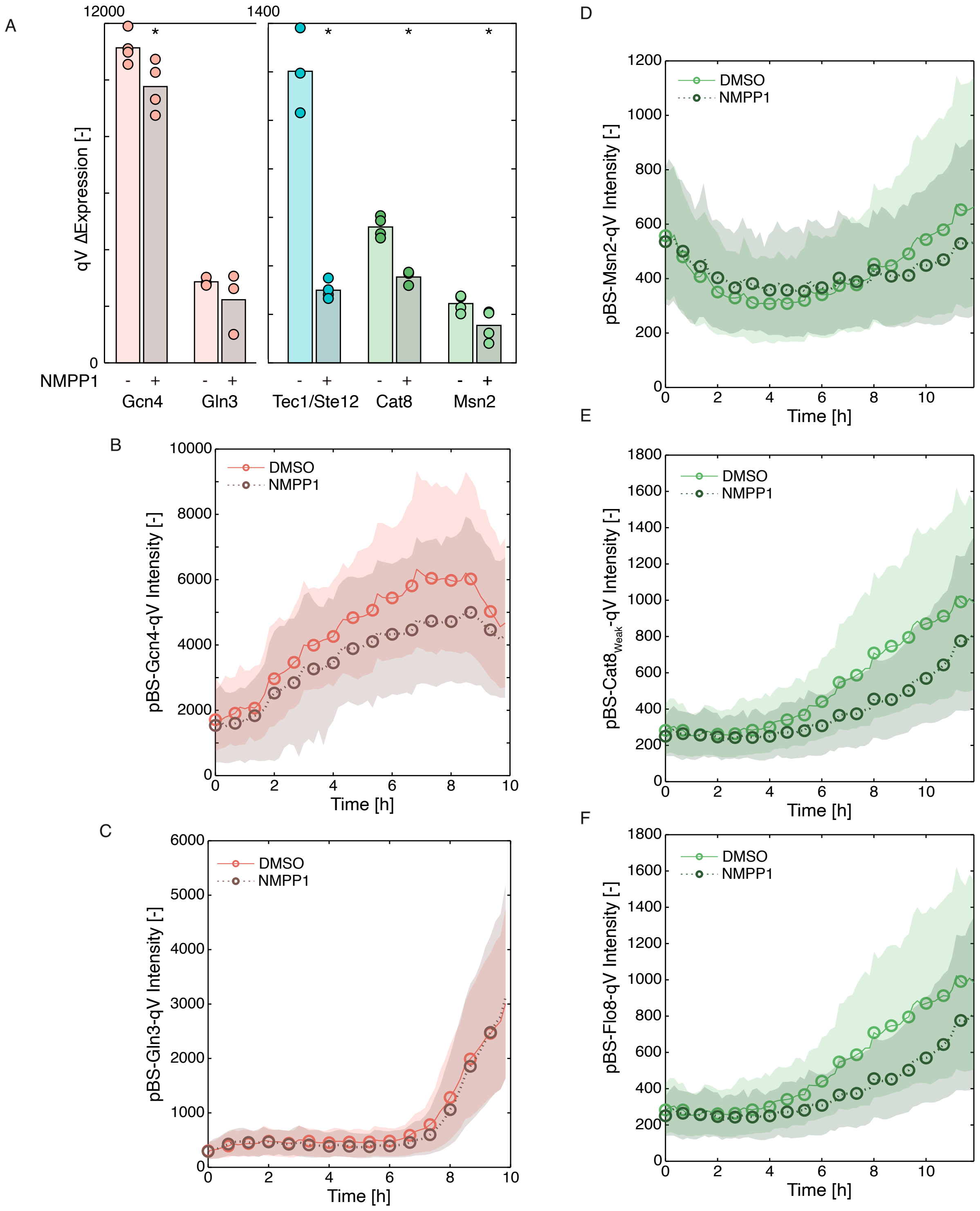
A
12000
1400
qV ΔExpression [-]
0
NMPP1
Gcn4
Gln3
Tec1/Ste12
Cat8
Msn2
B
DMSO
NMPP1
pBS-Gcn4-qV Intensity [-]
Time [h]
C
DMSO
NMPP1
pBS-Gln3-qV Intensity [-]
Time [h]
D
DMSO
NMPP1
pBS-Msn2-qV Intensity [-]
Time [h]
E
DMSO
NMPP1
pBS-Cat8Weak-qV Intensity [-]
Time [h]
F
DMSO
NMPP1
pBS-Flo8-qV Intensity [-]
Time [h]

Figure S8

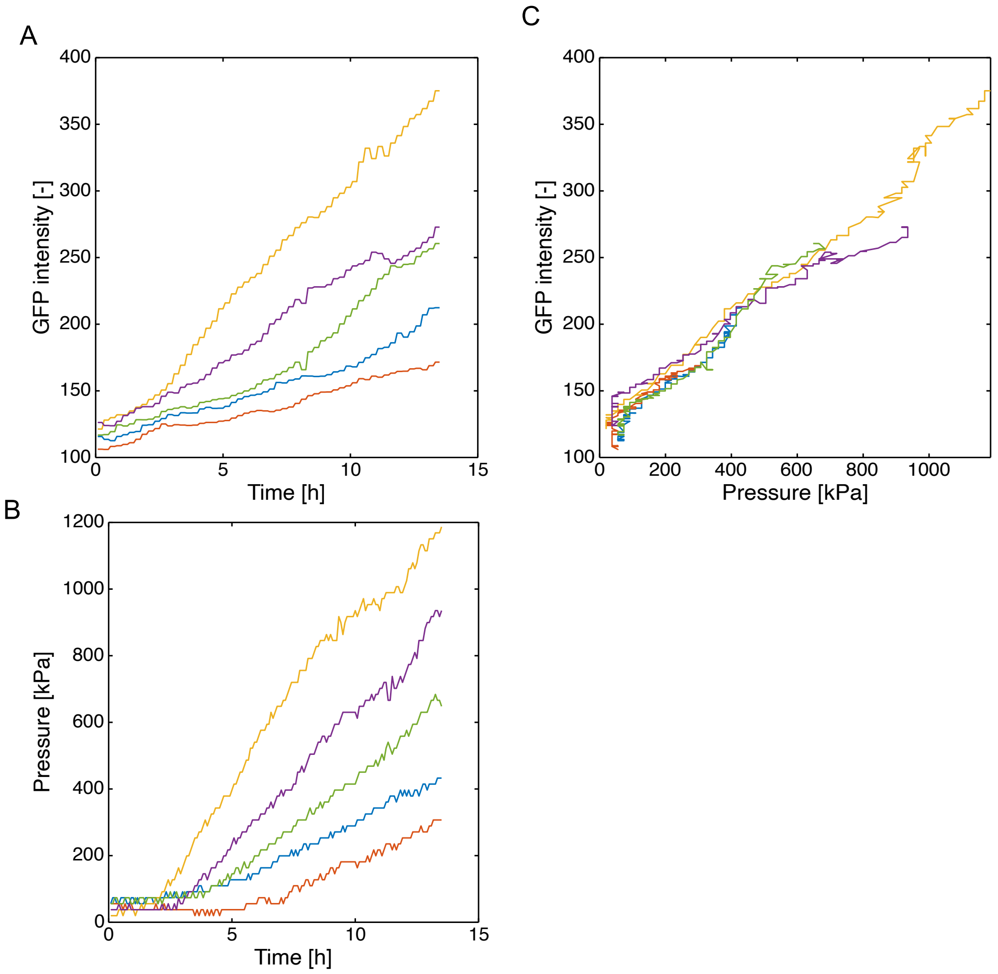

A
GFP intensity [-]
Time [h]
B
Pressure [kPa]
Time [h]
C
GFP intensity [-]
Pressure [kPa]


Figure S9
