## Supplementary figures and images for "Cellular integration of surface and nutrient signals during the early stage of filamentous growth in *Saccharomyces cerevisiae*"

### M1_pFLO11-qV_SLAD.gif

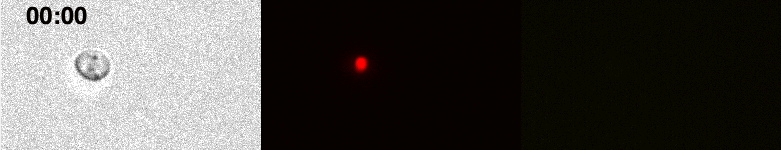

### M2_Flo11_endo.gif

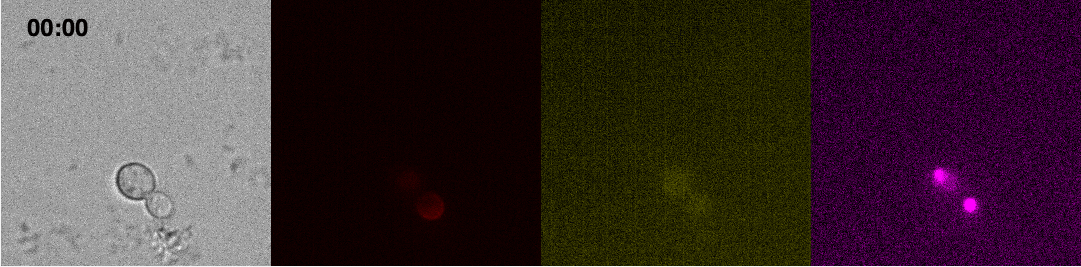

### M3_pYCT1-qV_SLAD.gif

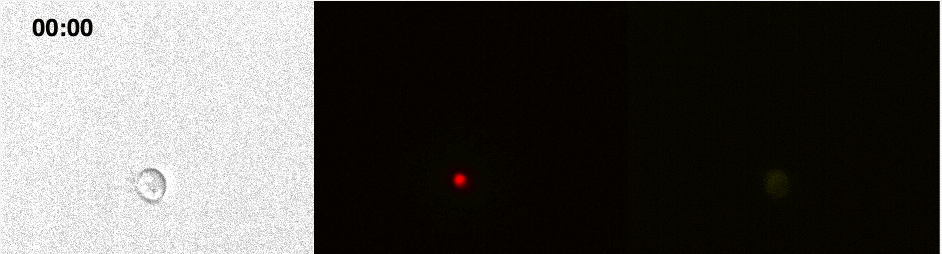

### M4_pBS_Ste12Tec1-qV_SAD.gif

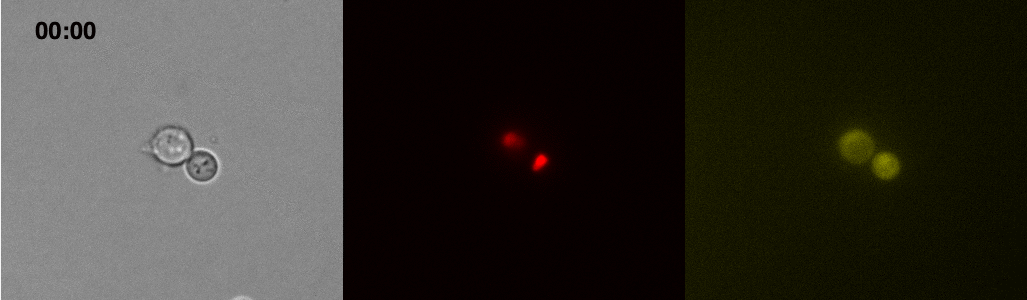

### M5_pBS_Ste12Tec1-qV_SALD.gif

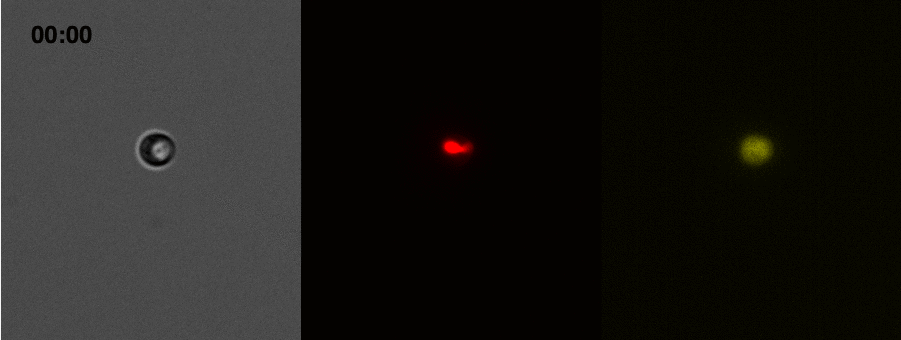

### M6_pBS_Ste12Tec1-qV_SLALD.gif

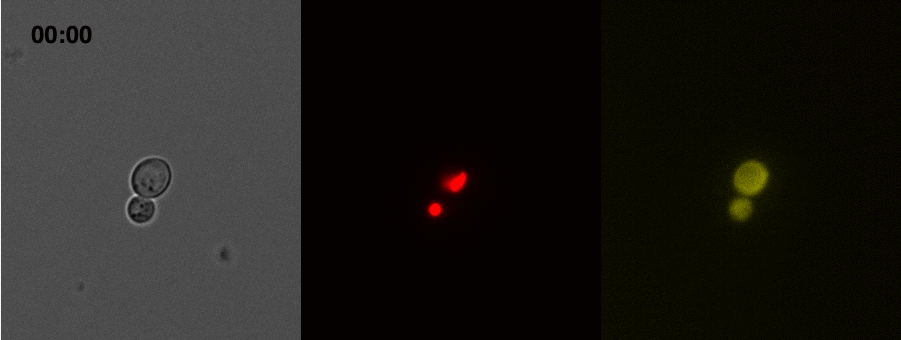

### M7_pBS_Ste12Tec1-qV_SLAD.gif

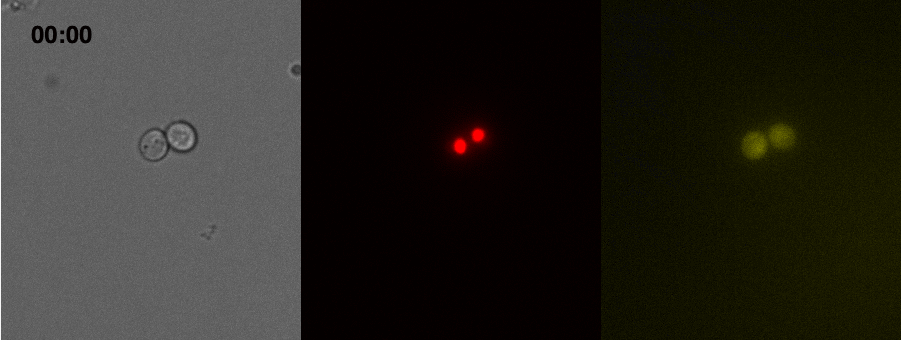

### M8_pBS_Gcn4-tSc_pBS_Gln3-qV_SLAD.gif

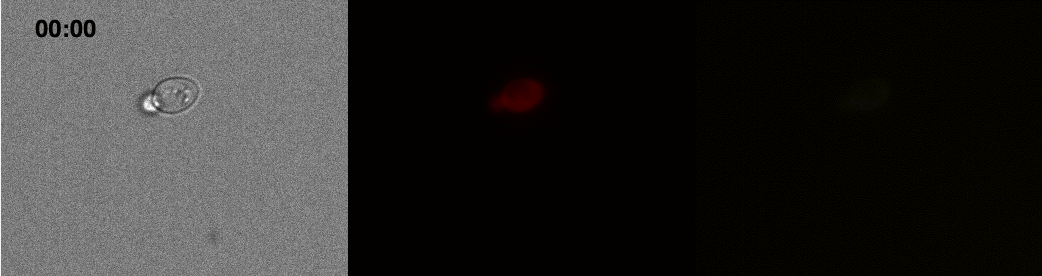

### M9_pBS_Ste12Tec1_Compress.gif

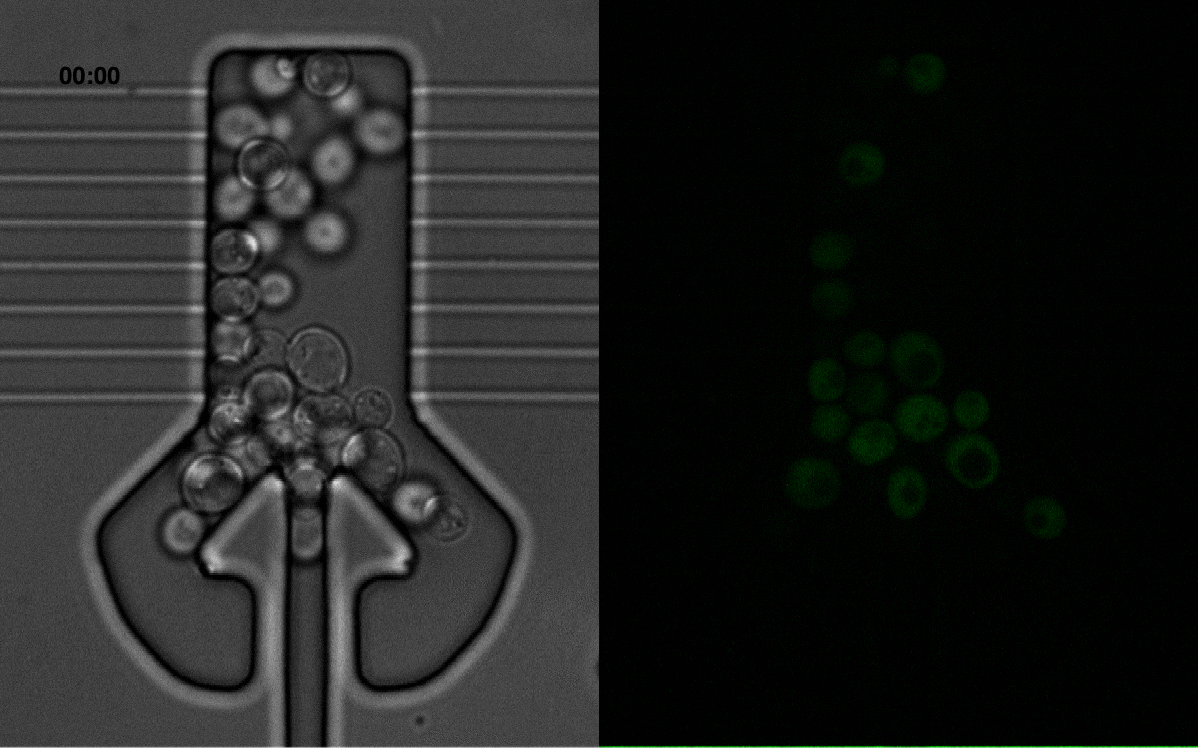

### M10_pNB_Ste12Tec1_Compress.gif

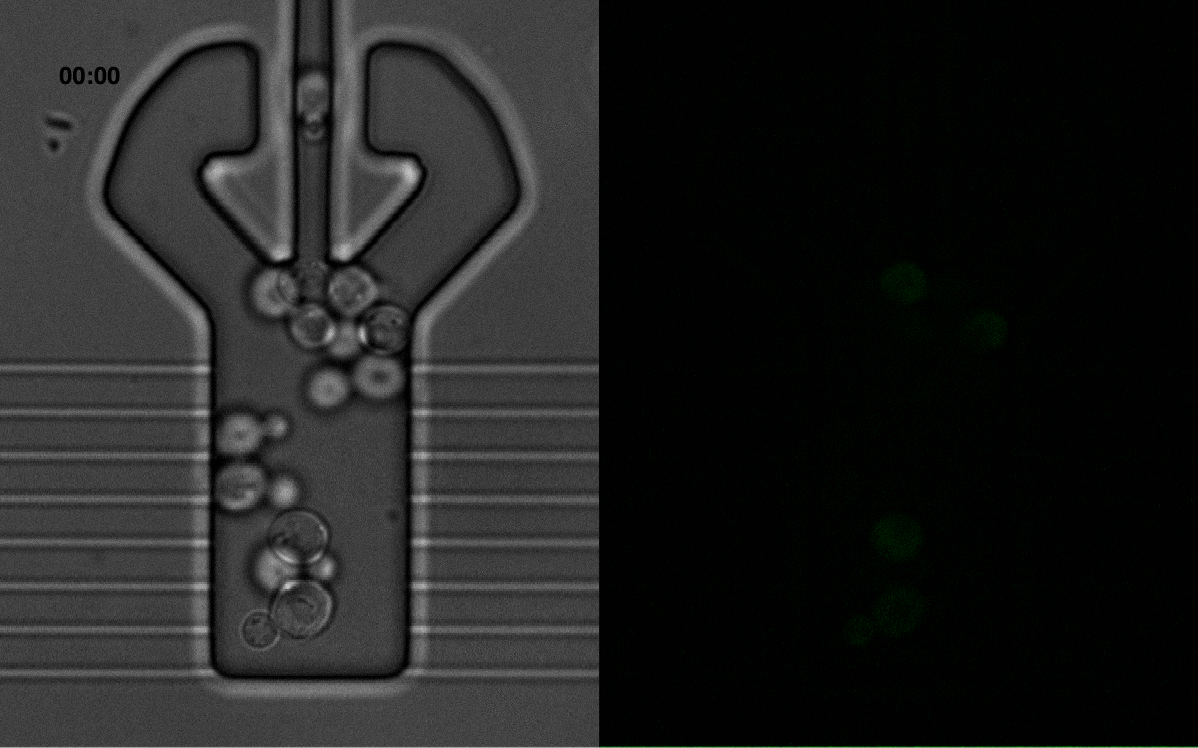
